# ER stress sensor Ire1 deploys a divergent transcriptional program in response to lipid bilayer stress

**DOI:** 10.1101/774133

**Authors:** Nurulain Ho, Haoxi Wu, Jiaming Xu, Jhee Hong Koh, Wei Sheng Yap, Wilson Wen Bin Goh, Shu Chen Chong, Stefan Taubert, Guillaume Thibault

## Abstract

Membrane integrity at the endoplasmic reticulum (ER) is tightly regulated and is implicated in metabolic diseases when compromised. Using an engineered sensor that exclusively activates the unfolded protein response (UPR) during aberrant ER membrane lipid composition, we identified pathways beyond lipid metabolism that are necessary to maintain ER integrity in yeast and are conserved in *C. elegans*. To systematically validate yeast mutants disrupting ER membrane homeostasis, we identified a lipid bilayer stress (LBS) sensing switch in the UPR transducer protein Ire1, located at the interface of the amphipathic and transmembrane helices. Furthermore, transcriptome and chromatin immunoprecipitation (ChIP) analyses pinpoint the UPR as a broad-spectrum compensatory pathway in which LBS and proteotoxic stress-induced UPR deploy divergent transcriptional programs. Together, these findings reveal the UPR program as the sum of two independent stress events and could be exploited for future therapeutic intervention.

## INTRODUCTION

The equilibrium between load of client proteins and protein folding is finely tuned within the endoplasmic reticulum (ER) environment, regulated by cellular pathways as part of the ER-protein quality control (ERQC). During periods of environmental stress [4], disease, infection or aging [5,6], ERQC is compromised, leading to the accumulation of misfolded proteins and causes ER stress. In turn, ER stress activates the evolutionarily conserved unfolded protein response (UPR) to restore ER homeostasis; if unresolved, chronic ER stress leads to apoptosis (for a review, see [7]). In metazoans, the three transmembrane protein transducers inositol requiring enzyme 1 (Ire1), protein kinase RNA (PKR)-like ER kinase (PERK) and activating transcription factor 6 (ATF6) sense ER stress and activate downstream cascades as part of the UPR program.

In addition, these transducers sense lipid bilayer stress (LBS) independently of the accumulation of misfolded protein in the ER lumen [2,8–12]. First identified as the sole ER stress transducer in *S. cerevisiae*, Ire1 is essential for cell viability during ER stress [13]. Ire1, through its downstream target mRNA and transcription factor *HAC1*, upregulates lipid biosynthetic genes [14]. Conversely, the deletion of lipid regulatory genes such as modulators of sphingolipid synthesis genes *ORM1* and *ORM2* in yeast or changes in the sphingolipids, dihydrosphingosine and dihydroceramide, in mammalian cells lead to lipid imbalance-induced UPR [11,15,16]. Similarly, increasing cellular saturated fatty acids through genetic manipulation or by exogenous supplementation strongly activates the UPR, likely triggered by a change in membrane fluidity [2,9,10,17,18]. Moreover, in obese mice and humans, altered membrane lipids composition in the liver and adipose tissues are associated with elevated UPR markers, further suggesting the ability of the UPR sensors to detect changes in membrane lipids [19,20]. Likewise, perturbing the levels of some membrane phospholipids, including phosphatidylcholine (PC), the most abundant phospholipid in the ER membrane [21,22], can lead to ER stress and UPR activation [10,12,19,23,24].

Despite the intimate relationship between lipid dysregulation and the UPR, our insight into how UPR transmembrane sensors such as Ire1 sense changes in the lipid composition of the ER membrane is incomplete. A form of yeast Ire1 (ΔIII Ire1), bearing a truncation in the LD, is still capable of activating the UPR through LBS by inositol depletion [8]. Moreover, in mammalian cells, a LD deletion Ire1 variant remained sensitive lipid perturbation at the membrane and induces the UPR [9]. Importantly, LBS activation of the UPR requires the integration of Ire1 transmembrane domain into the ER membrane [9], suggesting a direct sensing mechanism. A conserved amphipathic helix in proximity to the transmembrane helix of Ire1 drives the oligomerization of Ire1 during LBS [2]. However, mutating this amphipathic helix of Ire1 also severely diminished UPR activation during proteotoxic stress, suggesting that this mutation inactivates Ire1. These results point to a conserved UPR sensing and activation mechanism through LBS, that is independent of ER stress [25]. Altogether, these findings suggest the potential of specific domains within UPR sensors that monitor changes within the ER lipid bilayer to initiate an adaptive response.

In this study, we identified unexpected cellular perturbations inducing the UPR by LBS (termed as UPR^LBS^). These were found by monitoring the UPR activation in yeast in a genome-wide genetic screen of which the LD of Ire1 was found unnecessary for its activity. Several identified genes are conserved in metazoans, and their inactivation in *C. elegans*, similarly led to UPR^LBS^. In yeast, a strain lacking the *OPI3* gene was one of the strongest hits inducing the UPR. Since the homolog of *OPI3, Pemt*, is required for lipid homeostasis of membranes, we further characterized the activation mechanism of Ire1 in Δ*opi3* [26–28]. We identified a single residue within the interface of the Ire1 amphipathic and transmembrane helices that render it insensitive to LBS, when mutated, while retaining the capacity to activate the UPR by proteotoxic stress. Transcriptomic combined with ChIP-qPCR data revealed that the UPR program differs when activated by proteotoxic stress or LBS. Hac1 was found to associate to additional promoters of the genes *PIR3* and *PUT1* during LBS. Together, our data support a model where the UPR is a broad-spectrum compensatory pathway in which LBS and proteotoxic stress-induced UPR deploy divergent transcriptional programs.

## RESULTS

### A wide range of cellular perturbations activates Ire1 independently of its luminal domain

The activation of the UPR by Ire1 has been demonstrated to be specifically LBS-mediated by either depleting inositol, a phosphatidylinositol and sphingolipid precursor, or by supplementing saturated fatty acids [2,8,9]. Additionally, a selective screen of 17 knockout yeast strains was carried out to identify perturbations that are sensed by Ire1 containing a truncated LD (ΔIII Ire1) [8]. This study identified 9 genes with ER and metabolism related functions but fell short of providing a global picture of cellular perturbations that activate the UPR through Ire1 transmembrane domain. To address this gap, we generated a yeast Ire1 mutant lacking its entire LD (Ire1ΔLD) (Figure S1A). We exploited Ire1ΔLD inability to bind misfolded proteins and used it to monitor, on a global level, gene deletions that activate the UPR. These potential candidate genes could be involved in cellular processes that are necessary for ER membrane integrity. Using this tool, we carried out a genome-wide genetic screen in yeast to monitor the *in vivo* UPR activation with two query strains expressing either endogenous full length Ire1 and Ire1ΔLD (Figure 1A). These two query strains also express a green fluorescent protein (GFP) gene driven by the UPR element (UPRE)-containing promoter and a mCherry gene driven by a constitutive *TEF2* promoter [29]. The output was measured as the median of single-cell fluorescence ratio (GFP/mCherry) using high throughput flow cytometry.

**Figure 1.**
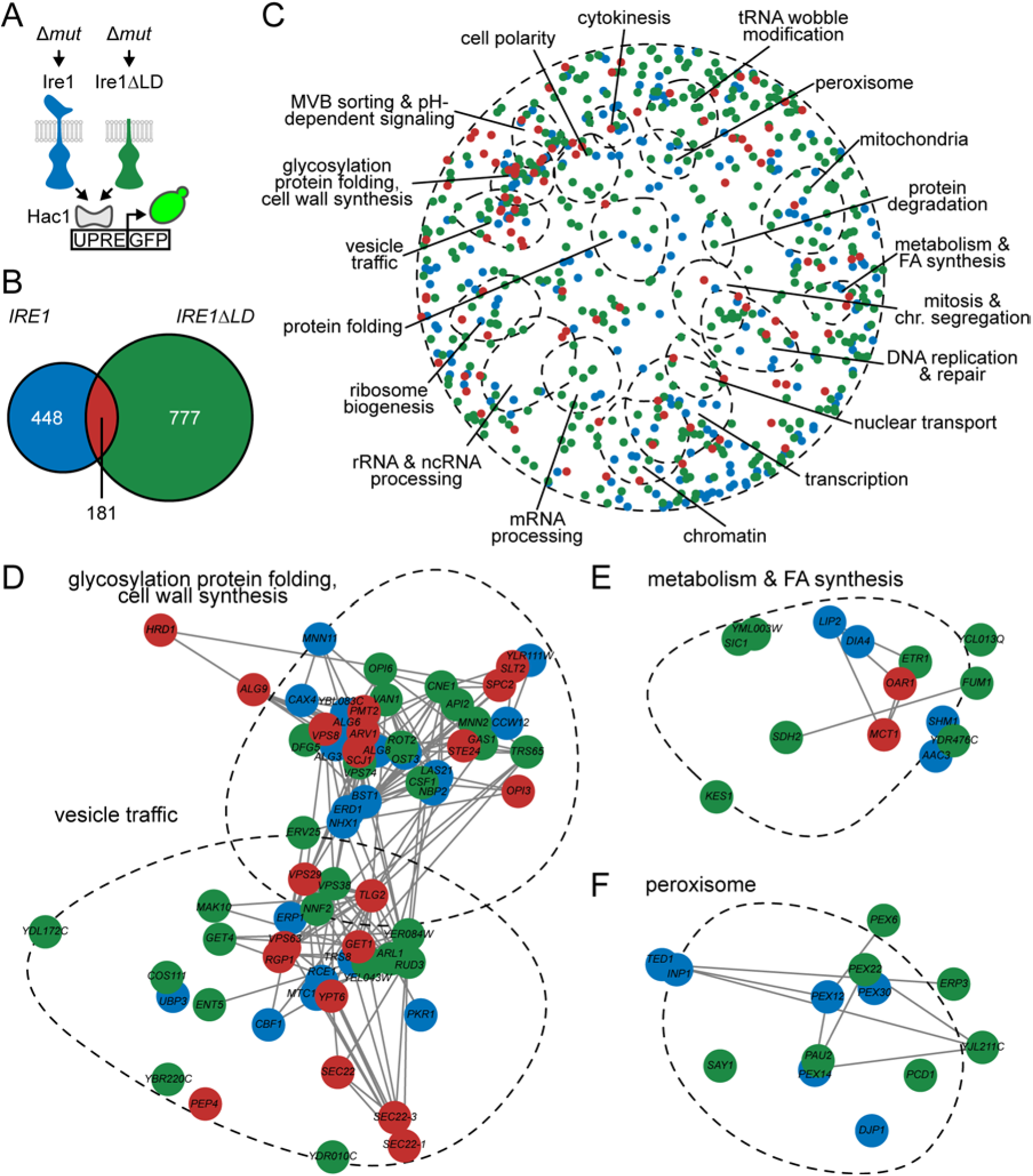
A high-throughput screen reveals gene clusters that mount a UPR response in *IRE1ΔLD* cells. **(A)** Strategy of the genome-wide high-throughput screen adapted from the synthetic genetic array methodology [1]. Query strains expressing *IRE1* or *IRE1*Δ*LD* were mated to the deletion strains. The resulting selected haploid strains were analysed by flow cytometry to measure the UPR activation (GFP) normalized to cytosolic mCherry. **(B)** Venn diagram depicting number of deletion mutants expressing *IRE1* (blue) or *IRE1ΔLD* (green). Overlap of *IRE1* and *IRE1ΔLD* deletion mutants are shown in red. Shown are number of strains giving fold changes that were ≥ 1.5 compared to the median GFP/mCherry signal. **(C)** Biological processes involving *IRE1* or *IRE1*Δ*LD* deletion mutants in the global cellular context [3]. Deletion mutants are color-coded as in (B). **(D-F)** Genetic interactions display genes associated to the ER involved in processes such as glycosylation, protein and cell wall synthesis, metabolism and fatty acid (FA) synthesis that are involved in UPRLBS, independent of peroxisome biogenesis. Deletion mutants are color-coded as in (B). See also Figure S1 and Table S1.

To functionally test our reporter system, the query strains were treated with proteotoxic stress inducing reagent dithiothreitol (DTT). The fluorescence signal ratio (GFP/mCherry) was significantly increased in full length Ire1 expressing cells but not Ire1ΔLD expressing cells (Figures S1B and S1C). This is indicative that DTT-induced UPR activation is dependent on the LD of Ire1. Moreover, we uncoupled the UPR activation by UPR^LBS^ from proteotoxic stress (termed as UPR^PT^) through the deletion of four genes previously identified to induce LBS [8]. As expected, deletion of these genes activated the UPR independent of the LD of Ire1 (Figures S1D and S1E). Interestingly, these four genes *STE24, SPC2, SCJ1* and *GET1* have roles in protein folding and translocation [30–33], providing the rationale that lipid composition could modulate both protein folding and/or protein trafficking through the ER. These genes are therefore required for proper ERQC as a result of UPR^LBS^. Together, these data demonstrate that both query strains are able to report UPR activation through proteotoxic stress and LBS. Both query strains endogenously expressing either full length Ire1 or Ire1ΔLD were subsequently mated with 4,847 strains from the *S. cerevisiae* deletion library [34] using the synthetic genetic array methodology [1] (Figure 1 and Table S1).

We identified 629 and 958 gene deletions that activated the UPR in an Ire1 LD dependent and independent manner, respectively (Figure 1B). Of note, the deletion of genes involved in the synthesis and transfer of N-linked glycosylation induced the UPR in query strain *IRE1* while no significant UPR activation was observed in query strain *IRE1*Δ*LD*. As N-linked glycosylation of proteins is necessary for the folding of nascent polypeptide in the ER [35], these hits further validate the screening of which query strain *IRE1*Δ*LD* insensitive to proteotoxic stress.

Other identified genes include dolichol-linked oligosaccharide synthesis genes *ALG3, ALG6*, and *ALG8* as well as oligosaccharyltransferase complex genes *OST3* (Table S1). Gene deletions that activated the UPR in Ire1ΔLD suggest these genes can activate UPR^LBS^ through Ire1 transmembrane domain. As previously reported [8], we identified that the ablation of genes *ARV1, ERV25, GET1, PMT2, OPI3, SCJ1, SPC2*, and *STE24* activated the UPR independently of Ire1 LD (Ire1^LD^), validating our approach (Figure 1D).

A total of 181 gene deletions were found to activate the UPR in both query strains. Most of these genes are closely related to the ER, suggesting that the lack of these genes specifically induce the UPR by disrupting processes related to ER membrane integrity (Figures 1C-1E, and Table S1). Several of these genes are implicated in UPR^LBS^ such as *TLG2*, required for endo-lysosomal fusion. The combinatorial deletion of *SEC14* and *TLG2* resulted in vesicular trafficking defects from the ER [36]. The *VPS* family of genes (*VPS8, VPS29, VPS61, VPS63*, and *VPS72*) involved in the endosomal to Golgi transport were similarly found to be required to maintain ER membrane integrity [37]. Another important cellular process found associated to UPR^LBS^ is the ER-associated protein degradation (ERAD) machinery. ERAD complex protein E3 ubiquitin ligase Hrd1 and its associated protein Hrd3 were found to be necessary for ER membrane homeostasis [38]. Hrd1 forms a ubiquitin-gated protein conducting channel for the retro-translocation of misfolded ER luminal protein across the ER lipid bilayer. Given that Hrd1-Hrd3 contacts the ER membrane bilayer, there is a potential for this complex to regulate changes to the membrane bilayer to buffer UPR^LBS^. This adds to the protective role of Hrd1 during UPR^PT^ through the degradation of misfolded proteins accumulated in the ER. Interestingly, peroxisome related genes did not activate the UPR in the overlap of query strains *IRE1* and *IRE1ΔLD* (Figure 1F). Together, these findings strongly argue that maintenance of vesicular trafficking and the ERAD pathways are necessary to maintain ER membrane integrity.

### Conserved cellular functions are necessary to maintain ER membrane integrity in *C. elegans*

To identify evolutionarily conserved cellular perturbations linked to UPR^LBS^, we carried out a reverse genetic RNA interference (RNAi) screen in the multicellular model organism *C. elegans*. We focused on identifying genes whose inactivation caused UPR activation through metabolic changes. Specifically, we depleted 1,247 genes predicted to be involved in metabolism (Figure 2A and Table S2) [39,40]; empty vector and mediator subunit 15 (*mdt-15*) RNAi clones served as negative and positive controls, as described [10]. To carry out the screen, synchronized stage 1 larvae (L1) bearing the IRE-1 activated and XBP-1 (*C. elegans* homologue of yeast Hac1) dependent reporter *hsp-4p::gfp* were subjected to RNAi, and GFP fluorescence was scored after 48 and 72 hours. The screen was completed in duplicate, and hits were subsequently confirmed in three independent validation experiments, yielding 36 RNAi clones that reproducibly induced *hsp-4p::gfp* fluorescence (Figure 2B and Table S2). In agreement with published data, we identified requirements for fatty acid desaturation enzymes *fat-4, fat-6*, and *fat-7*, PC synthesis enzymes *pcyt-1* and *sams-1*, the 3-hydroxy-3-methylglutaryl CoA synthase *hmgs-1*, and the Sarco-Endoplasmic Reticulum Calcium ATPase *sca-1*, validating our screen [10,12,41].

**Figure 2.**
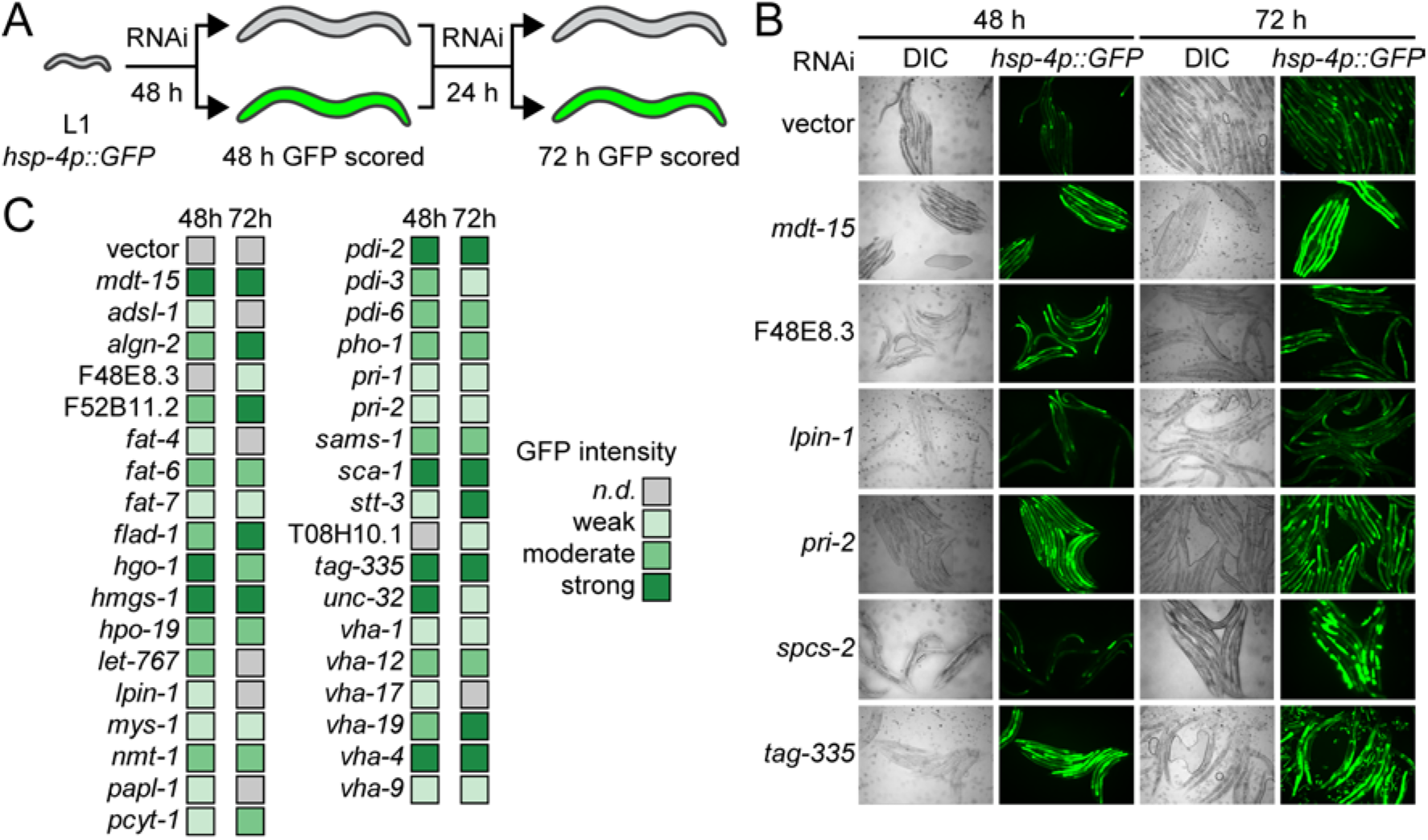
Membrane aberration activating the UPR is conserved in *C. elegans*. **(A)** Schematic of the *C. elegans* screen. Note that hits were scored as positive when above background fluorescence was detected at either 48 h or 72 h. **(B)** Representative fluorescence and DIC micrographs show *hsp-4p::gfp* worms at 48 h and 72 h after initiating growth on RNAi bacteria. Above background fluorescence indicates activation of the UPR. Vector refers to the empty RNAi vector negative control, *mdt-15* serves as positive control. **(C)** Summary of screen results. Shades of green indicate overall strength of *hsp-4p::gfp* induction as determined after three validation experiments. *n.d.*, non-detected. See also Tables S2 and S3.

We confirmed reliance on the canonical IRE-1 pathway by monitoring fluorescence after RNAi in a strain lacking XBP-1 (*xbp-1; hsp-4p::gfp*), and found that all 36 clones required *xbp-1* for induction (Table S2). In addition, we tested whether supplementation of choline, which can suppress UPR^LBS^ activation in worms defective for PC synthesis through the CDP-DAG pathway [10,12], is sufficient to suppress UPR activation. We observed partial rescue of RNAi clones *hmgs-1, lpin-1*, and *vha-4* as well as the expected complete rescue of *sams-1* RNAi treated animals, which are unable to synthesize PC through the CDP-DAG pathway (Table S2). Thus, 35 of 36 hits are likely inducing the UPR without dramatically altering PC levels.

We next tested whether the genes identified in the yeast screen are linked to UPR^LBS^ activation in *C. elegans*. Of 181 genes whose inactivation induced the UPR in both Ire1 and Ire1ΔLD yeast strains, we tested 38 and found that RNAi inactivation of one, the Signal Peptidase Complex Subunit homologue *spcs-2*, activated the *hsp-4p::gfp* reporter (Table S3). We also compared the 181 candidates from the screen done in yeast to the 36 candidates from our *C. elegans* screen to identify evolutionarily conserved processes or pathways whose impairment activates the UPR in both organisms. Some genes whose inactivation induced the UPR in *C. elegans* are essential in yeast (e.g. fatty acid desaturation genes *OLE1*, protein disulfide isomerase *PDI1*), preventing us from assessing conservation. Nevertheless, inactivation of genes in several pathways resulted in robust UPR induction across species, for example genes involved in PC synthesis, genes encoding the Vacuolar H^+^-ATPase, and several metabolic genes (Tables S2 and S3).

### Phospholipid perturbation activates Ire1 independently of its luminal domain

As a decrease in PC levels activated the UPR by Ire1ΔLD and IRE-1 in both yeast and *C. elegans*, we further characterized the activation mechanism of Ire1 in yeast cells lacking phosphatidylcholine biosynthesis gene *OPI3*, which displays a severe PC imbalance [23,24]. The disturbance of PC to phosphatidylethanolamine (PE) ratios in biological membranes is linked to ER stress and UPR activation in several model organisms [10,12,19,23,27]. Therefore, we utilized Δ*opi3* cells for a better understanding of PC-depletion sensing mechanism and downstream activated pathways by Ire1 as a means to develop future interventions.

To exclude the possibility of non-specific activation of the UPR due to the mislocalization of Ire1ΔLD, we assessed its subcellular localization by indirect immunofluorescence. Both Ire1-HA and Ire1ΔLD-HA colocalize with the ER marker Kar2 in both mutants (Figure S2A). We further validated the integration of Ire1ΔLD into the ER membrane by an alkaline carbonate extraction from the membrane fraction [42]. This method releases the contents of the membrane vesicles as well as peripheral proteins into the supernatant (S). Integral proteins remain embedded in the membrane and thus are found within the pellet fraction (P). Ire1 and Ire1ΔLD in both mutants were found in the pellet fractions together with the ER-localized transmembrane protein Sec61, indicating proper integration (Figure S2B).

To determine if Ire1^LD^ is required for UPR dependent survival of the synthetic lethal strain Δ*ire1*Δ*opi3*, growth assays were performed. As expected, both Δ*ire1* and Δ*ire1*Δ*opi3* strains expressing full-length *IRE1* grew in the presence of the proteotoxic stress inducer Tm (Figure 3A). Similar to Δ*ire1*, cells lacking *IRE1* LD failed to survive Tm-induced proteotoxic stress. In contrast, the expression of *IRE1ΔLD* was sufficient to rescue the synthetic lethality of Δ*ire1*Δ*opi3. IRE1*Δ*LD*Δ*opi3* cells displayed exacerbated growth rate during proteotoxic stress, potentially because these cells failed to further upregulate the UPR by the additional source of ER stress. We concluded that Ire1^LD^ is dispensable for restoring ER homeostasis through a functional UPR program during LBS.

**Figure 3.**
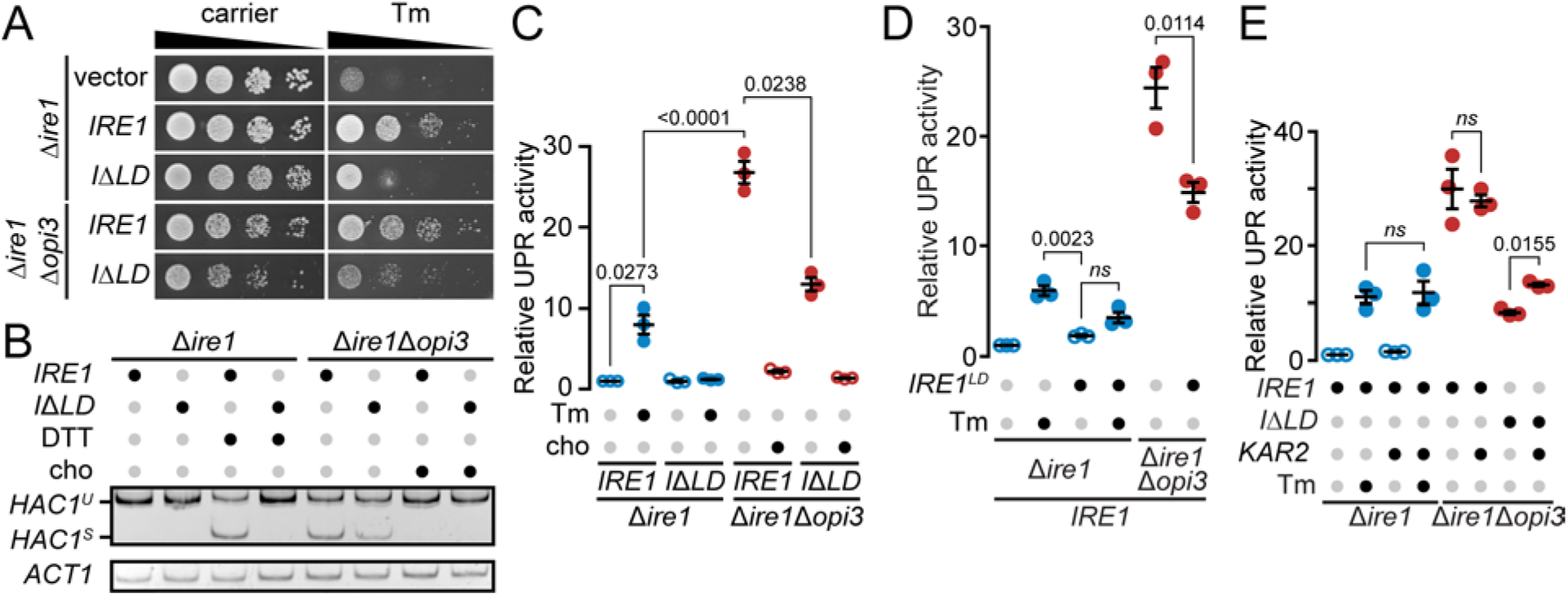
Ire1^LD^ is sufficient to uncouple the UPR activation triggered by LBS and proteotoxic stress in cells lacking *OPI3*. **(A)** The strains Δ*ire1* and Δ*ire1*Δ*opi3* expressing *IRE1* or *IRE1ΔLD* (*IΔLD*) were grown at 30°C and serial dilutions of the culture were spotted onto synthetic complete selective medium supplemented with 0.25 μg/ml tunicamycin (Tm), when indicated, and incubated until the appearance of colonies. **(B)** RT-PCR of unspliced *HAC1* (*HAC1^U^*) and spliced (*HAC1^S^*) mRNA. Media was supplemented with 1 mM choline (cho) or incubated 1 h with 2 mM DTT, when indicated. Actin (*ACT1*) was used as loading control. **(C-E)** UPR induction was measured using a β-galactosidase reporter assay of indicated strains **(C)** with the overexpression of *IRE1* luminal domain (*OE IRE1^LD^*) **(D)**, or *OE KAR2* **(E)**. Images shown are representatives of three independent experiments. Data shown is the mean ± SEM (n=3). Statistical analysis were subjected to paired two-tailed Student’s t-test. See also Figures S2 and S3.

To monitor the UPR activation, we assayed *HAC1* mRNA splicing under either proteotoxic stress or LBS. Under DTT-induced proteotoxic stress, *HAC1* mRNA was only spliced (*HAC1*^*S*^) in Δ*ire1* expressing *IRE1* while *IRE1ΔLD* failed to generate *HAC1*^*S*^ due to its inability in misfolded protein stress sensing (Figure 3B). Corroborating our growth assay results, during LBS, *IRE1ΔLD* was sufficient for *HAC1* mRNA splicing in Δ*ire1*Δ*opi3*. Choline supplementation inhibited *HAC1* splicing, presumably by alleviating LBS through the alternative PC synthesis CDP-choline pathway, suggesting that the UPR is specifically activated by a decrease in PC. To further validate our LBS-induced UPR model, we monitored the UPR activation using the UPRE-LacZ reporter assay [43]. As expected, Ire1^LD^ was necessary to induce the UPR during proteotoxic stress but dispensable during LBS (Figure 3C). It should be noted that the UPR activation in *IRE1ΔLD*Δ*opi3* was about half of Δ*opi3*, suggesting the strong UPR activation in Δ*opi3* is a combination of both proteotoxic- and LBS-induced ER stress, consistent with our growth assay (Figure 3A). In summary, our data validates that LBS directly activates Ire1 independently of its luminal domain to support cell survival.

To further uncouple the contribution of proteotoxic stress and LBS to the overall UPR program, we titrated the accumulation of misfolded proteins in the ER lumen by overexpressing (OE) *IRE1*^*LD*^ in the ER (Figure S2A) [44–46]. In WT cells, *OE IRE1*^*LD*^ was sufficient to significantly attenuate the UPR activation upon Tm treatment (Figure 3D). This result indicates that *OE IRE1*^*LD*^ is sufficient to prevent proteotoxic-induced UPR. Similarly, in Δ*opi3* cells, *OE IRE1*^*LD*^ significantly reduced the UPR activation by 39.1% compared to empty vector. The UPR attenuation is comparable to the decrease observed between Ire1 and Ire1ΔLD in Δ*opi3* cells (Figure 3C). These findings suggest that Ire1 senses both the accumulation of misfolded proteins and LBS resulting in a strong activation of the UPR in Δ*opi3* cells. Together, these findings support that solubilized *IRE1*^*LD*^ is capable of binding unfolded proteins in the ER lumen, partially inhibiting the UPR activation.

Next, we asked if the overexpression of the Hsp70 ER resident chaperone Kar2 (*OE KAR2*) attenuates proteotoxic stress-induced UPR as its overexpression mildly attenuates ER stress, possibly through the binding of misfolded proteins [47,48]. However, *OE KAR2* failed to reduce ER stress upon Tm treatment and during LBS with both Ire1 and Ire1ΔLD (Figure 3E). Unexpectedly, the UPR activity was significantly increased by *OE KAR2* in *IRE1*Δ*LD*Δ*opi3*, suggesting that the artificial expression level of this molecular chaperone might prevent timely substrate release during normal ER functions such as protein translocation, folding, and degradation [49–51]. More importantly, the inability of Kar2 to reduce ER stress reinforced the model by which Ire1^LD^ senses proteotoxic stress by direct binding to misfolded proteins and disfavor the dissociation of Kar2 from Ire1^LD^ to induce the UPR [47,52].

### LBS-activated Ire1 induces the UPR independently of its oligomeric state

An Ire1 variant missing its misfolded protein binding groove within the luminal domain, ΔIII Ire1, was previously reported to cluster into puncta upon inositol depletion in yeast [2,8], but with noticeable fewer puncta compared to proteotoxic stress, suggesting lower levels of dimerization by LBS. However, whether Ire1 lacking its entire LD is capable of oligomerization remains unknown [53]. To monitor Ire1 dimerization, we used a pair of split Venus fragments to monitor Ire1ΔLD dimerization *in vivo* by bimolecular fluorescence complementation assay (BiFC) [54], which localized to the ER (Figures S2D and S2E). As expected, co-expression of *IRE1-HA-VN173* and *IRE1-FLAG-VC155* produced fluorescent puncta at the ER upon DTT treatment, demonstrating Ire1 dimerization-dependent signal. (Figure 4A). Unexpectedly, in Δ*opi3* cells, fluorescent puncta were absent with both Ire1 and Ire1ΔLD while Ire1 was still responsive to DTT treatment in Δ*opi3*. To further assess this discrepancy with previous reports, LBS was induced with a 3 h inositol depletion in Δ*ire1* cells. Fluorescence puncta was absent in both Ire1 and Ire1ΔLD (Figure 4B and S2F). Next, to examine if tagged Ire1 variants are activated by LBS, we assayed *HAC1* mRNA splicing. Co-expression of split Venus *IRE1* fragments were sufficient to induce *HAC1*^*S*^ upon DTT treatment, inositol depletion and in Δ*opi3* cells while *HAC1*^*S*^ only accumulated in Δ*opi3* cells co-expressing split Venus *IRE1*Δ*LD* fragments (Figure 4C). Further validation of Ire1 dimerization in the Δ*get1*, Δ*scj1*, and Δ*ste24* mutants showed the absence of fluorescent puncta, consistent with the lack of dimerization observed in Δ*opi3* cells and inositol depleted cells (Figure 4D). As expected, Ire1 dimerization was evident in these mutants upon DTT treatment (Figure S2G). Together, these findings strongly argue that the formation of large Ire1 oligomers is mostly driven by proteotoxic stress whereas Ire1ΔLD is unable to form oligomers.

**Figure 4.**
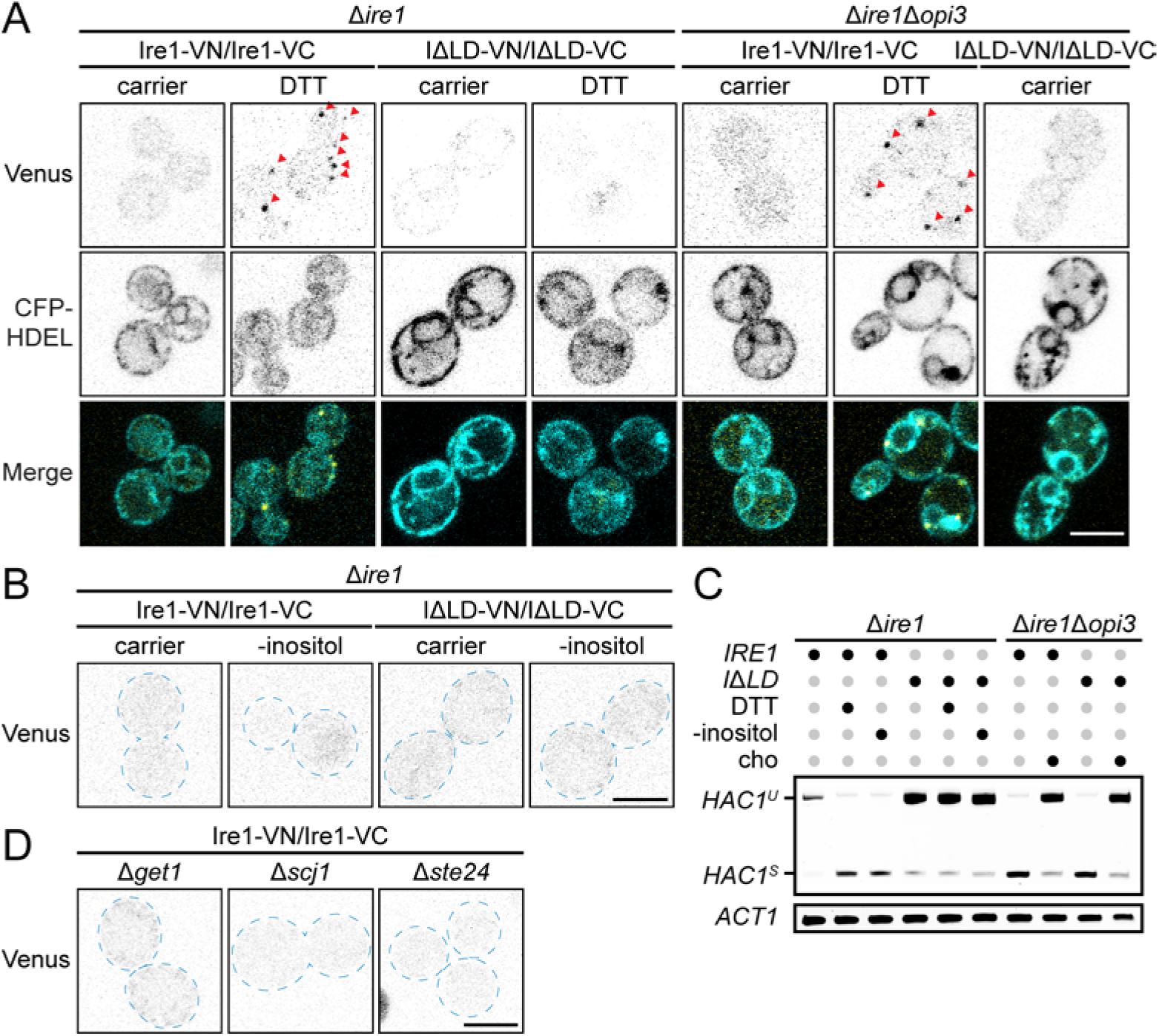
Ire1 forms higher ordered oligomers during proteotoxic stress that is absent during LBS. **(A-B)** Cells co-expressing the pair of split Venus fragments to monitor *IRE1-HA-VN173* and *IRE1-FLAG-VC155* or *IRE1ΔLD-HA-VN173* and *IRE1ΔLD-FLAG-VC155* to monitor dimerization *in vivo* by bimolecular fluorescence complementation assay (BiFC) in Δ*ire1* and Δ*ire1*Δ*opi3*. Cells were treated 1 h with 10 mM DTT **(A)** or depleted of inositol (-inositol) **(B)**, when indicated. CFP-HDEL ER was used as ER marker. **(C)** RT-PCR of unspliced *HAC1* (*HAC1^U^*) and spliced (*HAC1^S^*) mRNA. Media was depleted of inositol, supplemented with 1 mM choline (cho) or incubated 1 h with 2 mM DTT, when indicated. Actin (*ACT1*) was used as loading control. **(D)** Δ*get1*, Δ*scj1* and Δ*ste24* mutant co-expressing the pair of split Venus fragments and treated as in (A). Scale bar, 5 µm. Images shown are representatives of three independent experiments. Statistical analysis were subjected to paired two-tailed Student’s t-test. See also Figures S2 and S3.

Proper protein folding capacity in the ER lumen is important for cargo proteins to be adequately translocated into the ER lumen and properly modified so they can be correctly sorted to the final destination through COPII-coated vesicles [55]. Interestingly, several genes involved in COPII-coated vesicle transport to the Golgi apparatus and vesicular transport to the vacuole were identified to induce the UPR when lacking in the query strain *IRE1*Δ*LD* but not the *IRE1* strain, including *EMP46, ENT5, ERP2, ERV15*, and *ERV25* (Table S1) [56]. We previously reported that LBS-induced ER stress delays CPY translocation, with Ire1 facilitating this translocation by activating the UPR program [23]. We did not address whether the full UPR program is needed to facilitate this translocation. Therefore, we asked if cargo proteins are being accurately sorted and modified in cells bearing LBS-induced ER stress. CPY is a vacuolar enzyme that is processed in the ER lumen and sorted to the vacuole through the Golgi apparatus. We compared CPY translocation in cells expressing either full-length *IRE1* or *IRE1*Δ*LD* using a pulse-chase assay (Figure S3). CPY processing is delayed in cells expressing Ire1ΔLD under LBS-induced ER stress. Together with the genetic screen, the data suggest that the full UPR program is needed to alleviate the delay in cargo sorting and that Ire1^LD^ is necessary to maintain ER homeostasis if vesicle-mediated ER export of proteins is defective.

### A key arginine residue adjacent to the transmembrane domain of Ire1 is essential to sense LBS

A conserved amphipathic helix within Ire1^LD^, in proximity to the transmembrane helical domain, is necessary to drive Ire1 oligomerization upon inositol depletion in yeast [2]. Specifically, Ire1 mutants F531R and V535R are unable to induce the UPR to Ire1 WT levels upon both proteotoxic stress and LBS. As the introduction of the positive residue arginine within the amphipathic helix is likely to drastically disrupt the secondary structure of Ire1 and possibly decrease dimerization efficiency, we opted for a different approach to assess the role of Ire1 transmembrane helical domain in sensing LBS [57]. As expected, most of Ire1 transmembrane α-helix amino acid sequence is hydrophobic while two residues, within the core of the domain, are charged (Figure 5A). To maintain the hydrophobicity of the domain, we mutated the positive residue arginine into the polar uncharged residue glutamine at position 537 of yeast Ire1 [Ire1(R537Q)]. As expected, Ire1(R537Q) localized to the ER in both WT and Δ*opi3* cells (Figure S4). To validate the functionality of Ire1(R537Q) during proteotoxic stress, we carried out a spotting assay (Figure 5B). Both *IRE1(R537Q)* and *IRE1* exhibited similar growth on synthetic complete media supplemented with Tm. Next, we measured the splicing of *HAC1* mRNA by RT-PCR and qPCR (Figures 5C and 5D). During LBS in Δ*opi3* cells, no significant splicing of *HAC1* mRNA in *IRE1(R537Q)* was observed while *IRE1* WT strongly activated *HAC1* mRNA splicing. During proteotoxic stress, in cells treated with Tm, both Ire1 WT and R537Q spliced *HAC1* mRNA at similar levels. Remarkably, there was no significant splicing of *HAC1* mRNA by Ire1ΔLD(R537Q) during both proteotoxic stress and LBS.

**Figure 5.**
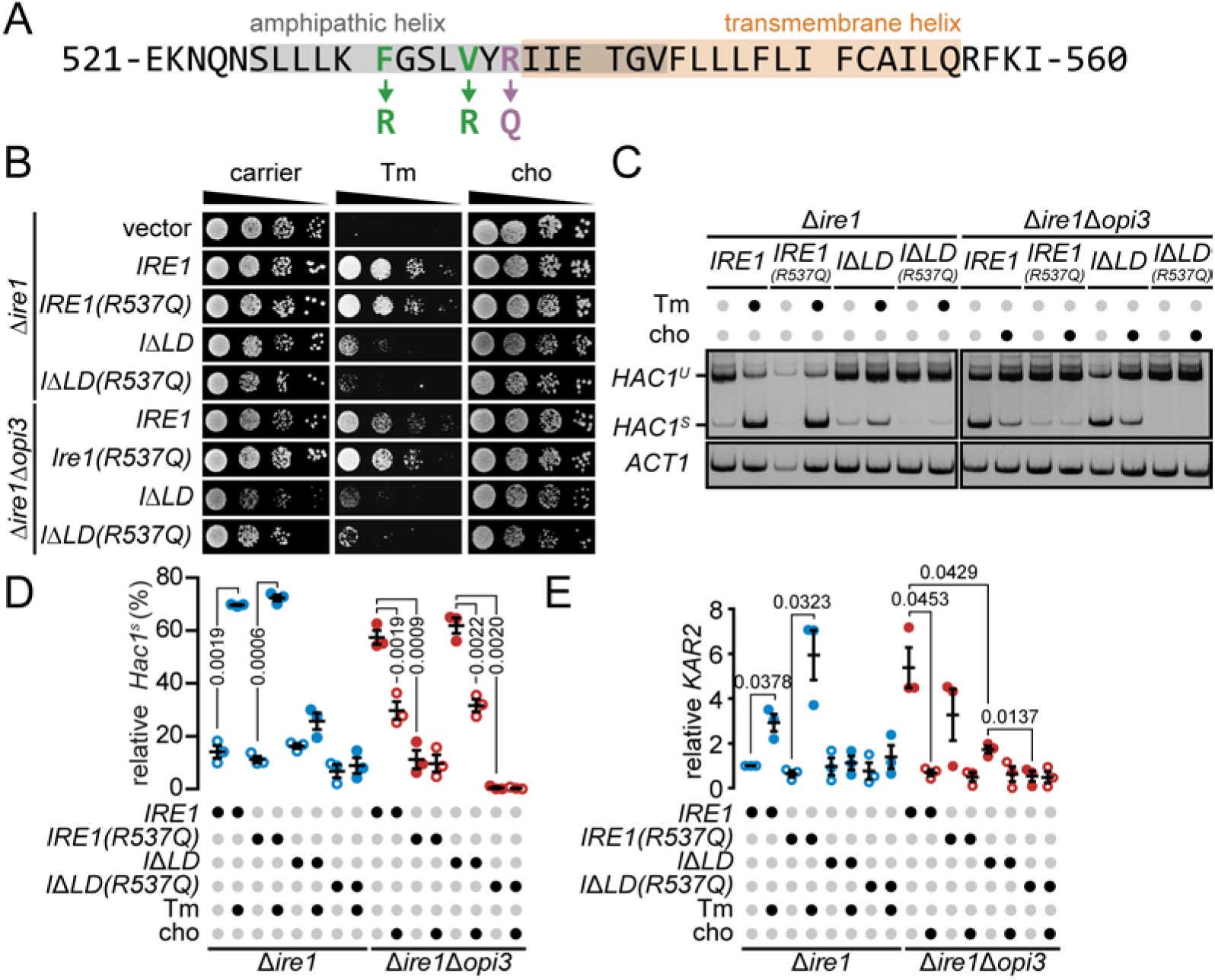
A key Ire1 arginine residue is critical to sense LBS. **(A)** Ire1 predicted amphipathic and transmembrane helices are highlighted in grey and orange, respectively. Point mutations highlighted in green were previously reported to be important in sensing both proteotoxic stress and LBS [2] and the point mutation in purple is part of this study. **(B)** The strains were grown, and serial dilutions of the culture were spotted onto synthetic complete selective medium supplemented with 0.25 μg/ml tunicamycin (Tm) or 1 mM choline (cho), when indicated, and incubated until the appearance of colonies. **(C)** RT-PCR of unspliced *HAC1* (*HAC1^U^*) and spliced (*HAC1^S^*) mRNA. Media was supplemented with 1 mM choline (cho) or incubated 1 h with 2 mM DTT, when indicated. Actin (*ACT1*) was used as loading control. **(D,E)** qPCR results comparing the splicing of *HAC1* (*HAC1^S^*) mRNA **(D)** or *KAR2* **(E)**. Media was supplemented with 1 mM choline (cho) or incubated 1 h with 2.5 µg/ml tunicamycin (Tm), when indicated. Images shown are representatives of three independent experiments. Data shown is the mean ± SEM (n=3). Statistical analysis were subjected to paired two-tailed Student’s t-test. See also Figure S4.

To further validate *IRE1(R537Q)* inability to induce *HAC1* mRNA splicing, we monitored the expression levels of *KAR2* mRNA which is induced through the Ire1-Hac1 axis upon ER stress (Figure 5E). As expected, *KAR2* was significantly upregulated upon Tm treatment only for full length Ire1 WT and R537Q. On the other hand, Ire1ΔLD(R537Q) failed to upregulate *KAR2* in Δ*opi3* cells, validating the requirement of the arginine residue within the transmembrane domain of Ire1 to sense LBS. Together, these findings are the first demonstration that Ire1 sensing of LBS can be disrupted while retaining the capacity to be activated by proteotoxic stress. Although the role of the R537 residue in sensing LBS is unlikely to be conserved in higher organisms, positive residue lysine is usually found at the edge of the transmembrane helical domain of IRE1 in metazoans, suggesting similar mechanistic uncoupling stresses in these proteins.

### A novel subset of genes is specifically upregulated by LBS-induced UPR

The UPR activates a broad-spectrum compensatory response during ER stress. To restore ER homeostasis, the required UPR-activated genes may vary depending on the source of stress [48,58]. To explore the deployed UPR program by LBS, DNA microarray analysis was performed in Δ*ire1* and Δ*ire1*Δ*opi3* cells expressing either *IRE1* or *IRE1*Δ*LD*. DTT-treated *IRE1* and *IRE1*Δ*LD* cells similarly upregulated 264 genes in comparison to unstressed WT cells. Therefore, these genes are modulated independently of Ire1^LD^, suggesting to be upregulated by other means than proteotoxic stress (Figures 6A, S5, and Table S4).

**Figure 6.**
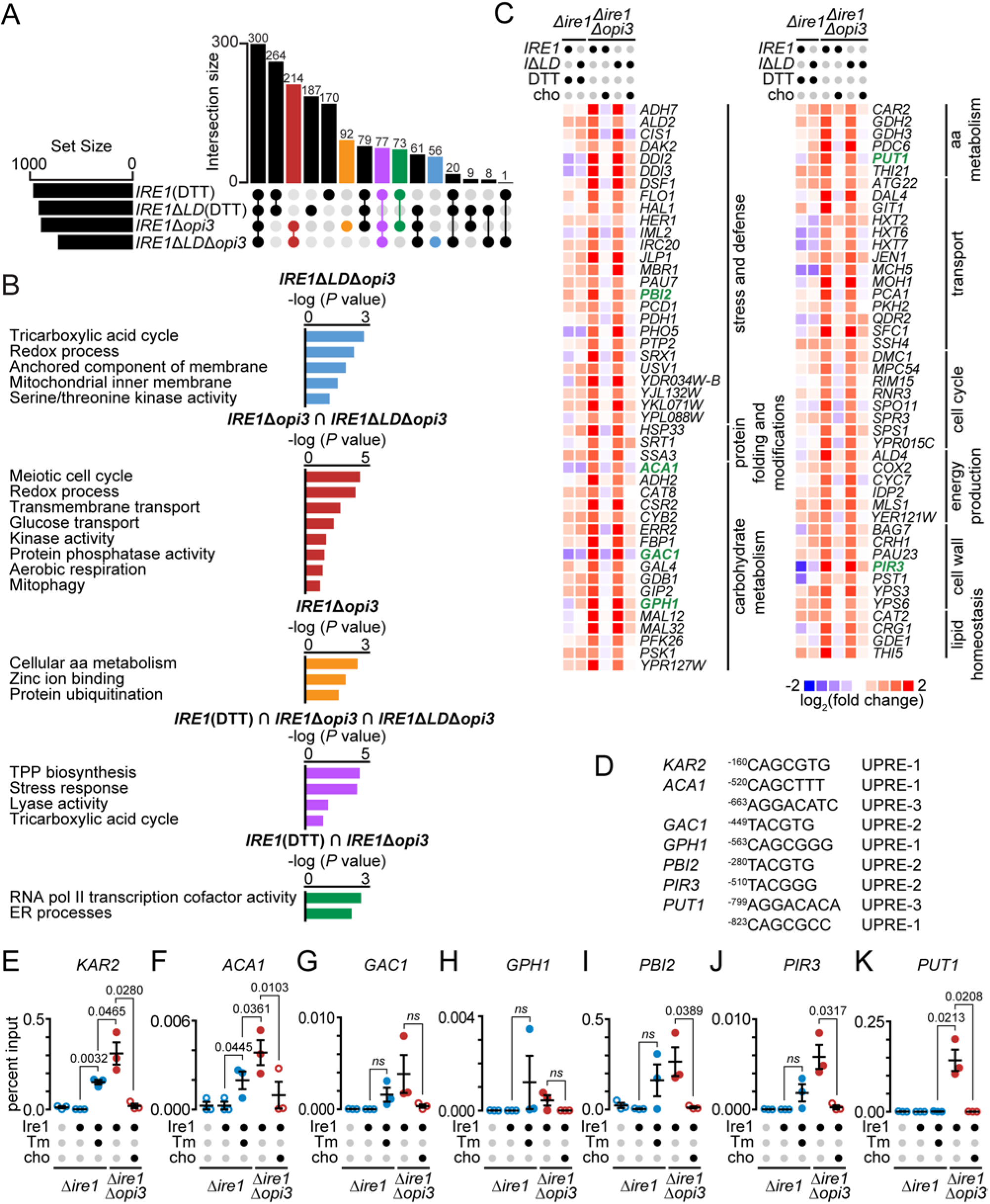
A subset of genes is upregulated by Hac1 specifically during LBS. **(A)** Diagram representing upregulated transcriptional targets of *IRE1* and *IRE1*Δ*LD* treated 1 h with 10 mM DTT, *IRE1*Δ*opi3*, and *IRE1*Δ*LD*Δ*opi3*. UpSet plot highlights intersections of selected group of genes differentially regulated during proteotoxic stress or LBS. Shown are number of genes giving fold changes that were >1.5 and at *P* < 0.05 (One-way ANOVA). **(B)** Bar plot of the GO analysis of genes upregulated in color-coded conditions as in (A). Genes are highlighted in yellow in Table S5. **(C)** Heat maps of selected LBS-induced genes. Based on log2 fold changes in gene expression normalized to untreated *IRE1* strain. Highlighted in blue are genes containing a predicted UPR element (UPRE) within the promoter region. **(D)** Potential Hac1 binding sites of different UPRE motifs within the promoter region of highlighted genes in (C). **(E-K)** ChIP-qPCR validation of predicted *HAC1* binding sites within the promoter regions of *KAR2* **(E)**, *ACA1* **(F)**, *GAC1* **(G)**, *GPH1* **(H)**, *PBI2* **(I)**, *PIR3* **(J)**, and *PUT1* **(K)**. *IRE1* were treated with 2.5 µg/ml of tunicamycin (Tm) and *IRE1*Δ*opi3* were supplemented with 1 mM choline (cho), when indicated. Data shown is the mean ± SEM (n=3). Statistical analysis were subjected to paired one-tailed Student’s t-test. See also Figure S5, Tables S4 and S5.

In Δ*opi3* cells, 214 genes were upregulated during LBS by both *IRE1* and *IRE1*Δ*LD* in comparison to unstressed WT cells. These genes were related to processes including transmembrane and glucose transport, aerobic respiration and mitophagy (Figure 6B). Interestingly, autophagic genes *ATG11* and *ATG32* required for ER-phagy were upregulated, suggesting a role of ER-phagy in buffering UPR^LBS^ [59]. However, a subset of these genes are likely to be upregulated independently of the UPR as we previously reported during PC depletion in yeast and *C. elegans* [12,23]. To identify genes specifically regulated through the UPR pathway during LBS, cells lacking *IRE1* or *HAC1* couldn’t be included as both are synthetic lethal in combination with Δ*opi3* cells [48,60]. Thus, we further analyzed the data to get around this hurdle. First, we performed functional annotation of upregulated genes in *IRE1*Δ*opi3* and/or *IRE1ΔLD*Δ*opi3*, and identified overlapping genes with full-length Ire1 expressing cells treated with DTT, using the gene ontology (GO) tool DAVID (Table S5). These 77 genes, in part, mount a response to stress pathways, such as previously identified UPR target *DER1* [61], validating the accuracy of the response of target genes (Figure 6B). Remodeling of the proteome and UPR activation occurs in Δ*opi3* mutants, and we confirmed the upregulation of known target genes such as secretory pathway members *SEC62* and *SEC72* in Δ*ire1*Δ*opi3* cells expressing Ire1 and Ire1ΔLD. Manual inspection of our microarray data highlights that known UPR target genes are strongly upregulated in *IRE1ΔLD*Δ*opi3*, in agreement with previous reports [12,14,23]. The exclusion of known UPR target genes within this cluster confirmed 139 genes were activated by LBS. Genes involved in ER stress and cellular stress resistance were strongly elevated as a global response to LBS. These include *SRX1* and *HSP33*, required for oxidative stress resistance [62], and *PDH1*, activated by diauxic shift [63]. Other genes involved in DNA replicative stress were upregulated, including glycogen degradation gene *GDB1*, carbohydrate metabolism gene *CAR2*, amino acid metabolism gene *PBI2*, and transporter gene *CYC7*.

Next, we analyzed genes that are differentially regulated between *IRE1ΔLD*Δ*opi3* and *IRE1*Δ*opi3* strains. This group of genes is potentially upregulated in response to UPR^PT^. For instance, the gene encoding the transporter protein *SEC24* was upregulated in a LD-dependent manner, suggesting upregulation only during UPR^PT^ [64]. *CIS1* which encodes for a protein required for autophagosome formation in *S. cerevisiae* is of particular interest because autophagy is required for achieving cellular homeostasis during the UPR [12].

To better understand how LBS affects UPR target genes, we further examined Hac1 specific targets within the positively upregulated 139 genes. As Hac1 binds to three known UPRE motifs [65], we performed bioinformatics analysis to identify putative UPRE consensus sequences in the promoters of the differentially regulated genes. We identified six genes containing the predicted UPRE-1, UPRE-2, and UPRE-3 motifs in their promoters that were upregulated by the UPR only during LBS (Figures 6C and 6D). The UPR only during LBS (Figures 6C and 6D). The upregulated genes include stress and defense gene *PBI2*, carbohydrate metabolism genes *ACA1, GAC1* and *GPH1*, amino acid metabolism gene *PUT1*, and cell wall synthesis gene *PIR3*. These findings suggest that Hac1 promotes metabolic processes (amino acid and carbohydrate metabolism) for the restoration of cellular homeostasis to compensate for the lack of proper lipid biosynthesis during LBS. Similarly, genes maintaining cell wall integrity were upregulated, arguing that cell wall stress occurs during LBS-induced ER stress and pinpointing to the importance of coordinating cell wall biogenesis and the UPR response.

To validate the putative LBS-specific UPREs, we performed chromatin immunoprecipitation (ChIP) in Δ*ire1* mutants expressing HA-Hac1 and Ire1. Immunoprecipitation specific to HA-Hac1 was used to pull down associating genes in the presence of protein stress and LBS. Primers were designed approximately 50 bases upstream and downstream of the putative UPRE for quantitative analysis of these genes. *KAR2* was incorporated to the ChIP assay as it contains a well characterized UPRE-1 within its promoter region (Figure 6E) [65]. Hac1 association to KAR2 promoter was indeed increased during Tm treatment. During Tm-induced protein stress, qPCR data confirms the absence of fold enrichment of the five gene promoters *GAC1, GPH1, PBI2, PIR3*, and *PUT1* (Figures 6G to 6K). *ACA1* promoter was significantly enriched by Hac1 binding during protein stress (Figure 6F). *ACA1* belongs to the during protein stress (Figure 6F). *ACA1* belongs to the family of bZIP proteins (including Hac1) and acts as ATF/CREB activators [66]. Contrary to what has been reported, *ACA1* does not bind the UPRE and is not involved in the unfolded protein response. Our findings, however, suggest that *ACA1* is a UPR target gene, possible through carbon source regulation [66]. In contrast, Hac1 was found to be associated to the promoters of PBI2, PIR3 and PUT1 during LBS, validating our transcriptomic data where these genes are upregulated by UPR^LBS^. These data strongly argue that the UPR transcriptional program is an adaptive response adjusted to the source of stress.

## DISCUSSION

In the last decade, it emerged that yeast Ire1 contains another independent sensing domain to monitor LBS at the ER [2,8]. Conserved in higher eukaryotes, IRE1 is similarly activated by LBS together with PERK [9] while ATF6 is activated by an increased level of sphingolipid species [11]. Recently, a detailed LBS-sensing mechanism of yeast Ire1 through its amphipathic helix revealed rotational orientations that stabilized its activation during proteostatic and lipostatic ER stress [2]. As LBS was mostly introduced by inositol depletion or saturated fatty acid excess in yeast and mammalian cells, the breath of cellular perturbations sensed by Ire1ΔLD still remained unclear. To address this knowledge gap, we carried out a genome-wide high throughput genetic screen and identified a subset of genes with various functions necessary to maintain ER membrane integrity in yeast and *C. elegans* (Figures 1–4). Furthermore, we identified an essential residue at the interface of the amphipathic and transmembrane helices of Ire1 that senses ER membrane integrity while being dispensable to activate the UPR by proteotoxic stress (Figure 5). By uncoupling LBS- and proteotoxic-induced UPR, we demonstrated that the UPR program is a broad-spectrum compensatory pathway with divergent transcriptomes (Figure 6).

Conical PE and cylindrical PC promote negative and minimal membrane curvature, respectively [7,67,68]. The phospholipid intermediate *N*-monomethyl phosphatidylethanolamine (MMPE), generated during *de novo* synthesis of PC from PE, exhibits physical properties similar to PE which becomes highly abundant under the ablation of *OPI3* [23,24]. The combination of a virtual absence of sterol at the ER and the replacement of PC with MMPE, both contribute to the stiffening of the membrane [24,69–72]. Consequently, it is conceivable that during LBS alone, the binding affinity of Ire1 is insufficient to counteract the stiffening of the membrane in forming homodimers or higher oligomers. In contrast, the large accumulation of unfolded proteins in the ER, by Tm or DTT, might be necessary to promote Ire1 dimerization through the binding of Ire1 luminal domain during LBS [47,53,73,74]. The arginine residue located at the interface of the amphipathic and transmembrane helices of Ire1 is essential for LBS sensing, suggesting a conformational change inducing Ire1 activation (Figure 5).

In agreement with our findings, no Ire1α homodimers or oligomers were detected by palmitic acid-induced LBS [18]. Accordingly, non-oligomerized Ire1 spliced *HAC1* mRNA during PC or inositol depletion (Figures 4C and 4D). Although reports have shown that a dimerization-dependent conformational switch is required to activate Ire1 RNase domain, Ire1 RNase activity is preserved as a monomer [75,76]. Kinase-inactive Ire1 splices *HAC1* mRNA while displaying defects in Ire1 deactivation [75], leading to chronic ER stress. On the other hand, ER-stress induced clustering of low endogenous Ire1 abundance [77] might be insufficient to be detected by conventional methods during LBS. For instance, the reported LBS-induced Ire1 puncta are noticeably weaker and less abundant when compared to puncta of proteotoxic stress, suggesting higher Ire1 oligomerization states during proteotoxic stress [2,8]. Similar to our findings, residues W457 and S450 of the transmembrane domain of mammalian Ire1α are required for stable dimer formation during membrane saturation by palmitic acid [78]. Together with our findings, it suggests that Ire1 activation mechanism differs between proteotoxic stress and LBS. These two activation routes might work synergistically or independently to transduce the UPR signal (Figure 7).

**Figure 7.**
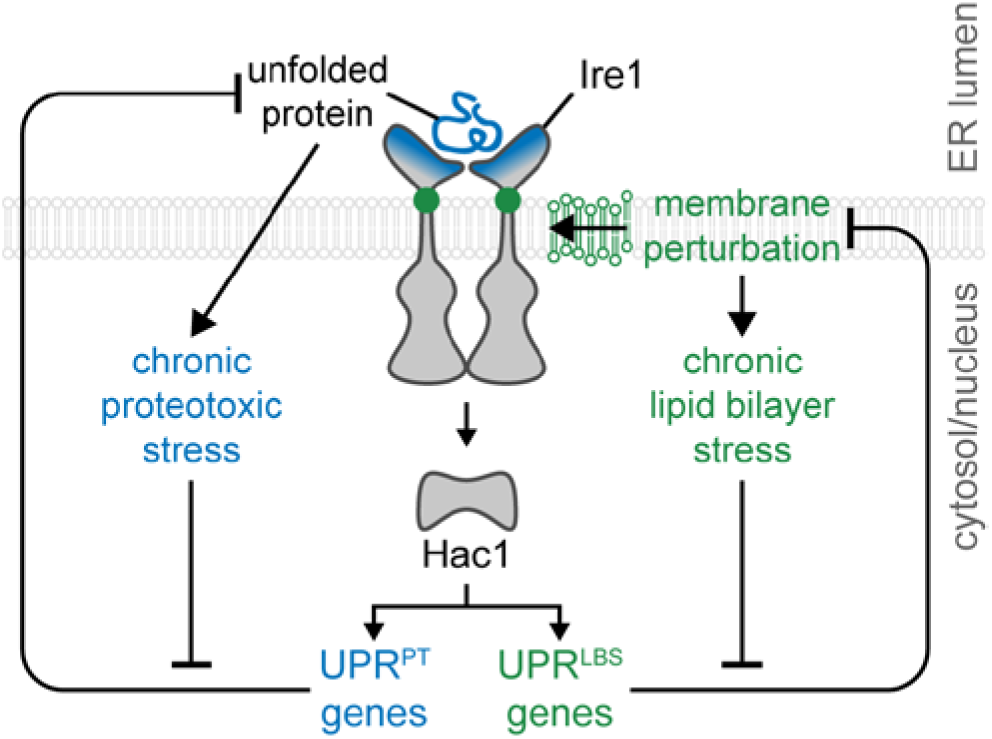
ER stress sensors induce differential UPR programs to restore ER homeostasis. Upon the accumulation of unfolded proteins or during membrane perturbation, ER stress sensor Ire1 activates downstream transcription factors resulting in the upregulation of UPR target genes. Initially a linear stress response pathway, the UPR transcriptional program diverges in response to unresolved ER stress – UPR^PT^ and UPR^LBS^. The UPR program might become more targeted to a specific need if the source of stress is constitutive or chronic such as in the context of diseases.

To overcome ER stress, the UPR broadly shifts the transcriptome along with differential expression levels across cell types, providing cellular robustness [48,79,80]. Once Ire1 senses ER stress, the Hac1-mediated response is assumed to be linear until ER homeostasis is restored. However, there are several lines of evidence suggesting the contrary in yeast and mammals, particularly during LBS. Previously, we demonstrated that the UPR transcriptional program is modulated through differential target gene expression depending on the source of stress [48]. The resulting differential UPR transcriptional program included the ablation of Hsp70 co-chaperone *SCJ1* which was identified to induce UPR^LBS^ in our genome-wide genetic screen (Figures 1 and S1). In *C. elegans*, we recently demonstrated that the transcriptome diverges dramatically between UPR^PT^ and UPR^LBS^, including hundreds of genes upregulated in an IRE-1-dependent (IRE1α homologue) manner during LBS [12]. Similarly, in mammalian cells, the ATF6-modulated UPR program diverged upon LBS in comparison to proteotoxic stress [11]. In agreement with these findings, *HAC1* mRNA level was revealed to be enhanced by a bipartite signal, misfolded proteins and either inositol depletion or temperature shift [61]. In the presence of one of the latter signals in Δ*ire1* cells, the levels of *HAC1* mRNA more than doubled, indicating an Ire1-independent mechanism maintaining protein quality control, mounting an alternative transcriptional program. This higher magnitude of UPR termed as the “Super-UPR”, could be classified with its own transcriptional program.

Additionally, another group demonstrated the autoregulation of *HAC1* during period of extreme and prolonged ER stress by a positive feedback loop of Hac1 binding to its promoter [81]. There is a delicate balance faced by Ire1 in responding to the stress it encounters and a finely tuned response for the activation of particular genes to adapt to cellular changes [61]. In addition to transcriptional regulation, transcription factor Hac1 is regulated by multiple factors at the protein level. Gcn4, a transcriptional activator of amino acid biosynthetic genes, works synergistically with Hac1 at the promoter of UPR target genes [82]. In accordance with these findings, the identification and validation of three genes exclusively upregulated during UPR^LBS^ supports the integration of multiple cellular stimuli to mount a divergent transcriptional response by Hac1 (Figure 6). The functional relevance of each of these three validated genes, *PBI2, PIR3*, and *PUT1*, in buffering lipid bilayer stress still remains to be elucidated. Additionally, a large proportion of the characterized 381 genes, upregulated by UPR^PT^ in yeast [14,23,83], lack one of the three UPR elements. Possibly, additional unidentified *trans*-acting factors regulate Hac1 transcriptional factor [82].

The UPR is associated with numerous physiological processes in addition to protein quality control [84]. Several diseases including diabetes, non-alcoholic fatty liver disease (NAFLD), liver failure, cystic fibrosis, Alzheimer’s disease, Huntington’s disease, Parkinson’s disease, and cancer are linked to the UPR pathway [85,86]. Despite mounting evidences on the role of the UPR in metabolic diseases, the contribution of the ER membrane composition in activating the UPR is poorly understood. Our genome-wide genetic screen revealed a wide variety of cellular processes that are necessary to maintain ER membrane integrity (Figure 1). For instance, UPR^LBS^ was predominantly induced by genetically disrupting cellular pathways related to fatty acid and phospholipid biosynthesis, vesicular trafficking, and ER-phagy. Intuitively, low PC- and palmitic acid-induced LBS disrupt ER structure and integrity of which the UPR transcriptional program is essential for cell survival [23,87]. In accordance with our findings, the selective autophagy pathway ER-phagy is required to maintain cellular homeostasis by recycling ER membrane during ER stress [80,88]. A similar requirement of vesicular transport from the ER might be necessary to remove the otherwise excess of ER membrane. Although lipid synthesis is tightly regulated according to cellular needs, the overall buffering of ER membrane integrity is undoubtedly the coordinated effort of multiple regulatory pathways.

In this report, we show clear evidence linking the UPR to ER membrane integrity which implicates pathways beyond lipid metabolism. To overcome LBS, the activation mechanism of Ire1 by sensing fluctuation at the ER membrane diverges greatly to the well-studied activation mechanism during proteotoxic stress. In addition, through a mechanism that remains unclear, transcriptional factor Hac1 deploys a curated UPR transcriptional program to restore cellular homeostasis during LBS. Taken together, the data demonstrate the remarkable diverse cellular pathways working in concert with the UPR to maintain ER membrane integrity. How each of the regulatory pathways contribute to UPR-associated metabolic diseases will be the challenge of future studies.

## MATERIAL AND METHODS

### Strains and antibodies

*S. cerevisiae* strains used in this study are listed in Table S6. Strains were prepared using standard transformation protocols. Anti-HA mouse monoclonal antibodies HA.11 (Covance, Princeton, NJ), anti-FLAG mouse M2 monoclonal antibody (Sigma-Aldrich, St. Louis, MO), Anti-Kar2 rabbit polyclonal antibody, anti-CPY rabbit polyclonal antibodies, and anti-Sec61 rabbit polyclonal antibody were gifts from Davis Ng (Temasek Life Sciences Laboratories, Singapore). Secondary antibodies goat anti-mouse IgG-DyLight 488 (Thermo Fisher, Waltham, MA), goat anti-rabbit IgG DyLight 550 (Thermo Fisher, Waltham, MA), goat anti-mouse IgG-IRDye 800 (LI-COR Biosciences), and goat anti-rabbit IgG-IRDye 680 (LI-COR Biosciences, Lincoln, NE) were commercially purchased.

### *C. elegans* strains and RNAi constructs

All strains were grown at 20°C using standard *C. elegans* methods, as previously described [89,90]. Nematode growth medium (NGM) agar plates were seeded with *E. coli* strain OP50 for normal growth or with HT115 bacteria for RNAi feeding, as indicated. The wild-type N2 Bristol, *atf-6(ok551) X, ire-1(ok799), pek-1(ok275) X*, SJ4005 (*hsp-4p::gfp*), and SJ17 (*xbp-1(zc12) III;zcIs4 [hsp-4p::GFP] V*) strains were obtained from Caenorhabditis Genetic Center (CGC). RNAi was performed using solid NGM-RNAi media, i.e. NGM containing 25 μg/mL carbenicillin, 1 mM isopropyl β-D-1-thiogalactopyranoside (IPTG), and 12.5 μg/mL tetracycline, seeded with the appropriate HT115 RNAi bacteria. All RNAi clones including positive controls *mdt-15* were from the Ahringer library and were sequenced prior to use.

### Plasmids used in this study

Plasmids and oligonucleotides used in this study are listed in Tables S7 and S8, respectively. Plasmids were constructed either by restriction or Gibson cloning. All coding sequences of plasmid constructs used in this study were fully sequenced. The plasmid pGT0421 containing HA tagged *HAC1* in pMR366 was constructed as previously described [48]. The plasmid pGT0330 containing endogenously *IRE1* was generated as previously described [91]. The plasmid pGT0201 containing *IRE1ΔLD* was generated by amplifying the endogenous promoter and signal sequence (fragment 1) and transmembrane cytosolic domain (fragment 2) with primer pairs GTO275-276 and GTO277-GTO278 respectively, from template WT genomic DNA (WT gDNA). The fragments were further digested with *XhoI* and *PstI* (fragment 1) and *Pstl* and *Notl* (fragment 2) before ligation into a pRS313 *XhoI* and *NotI* linearized plasmid. The plasmids pGT0223 and pGT0225 were generated by WT gDNA amplification using primer pair HWO15-16 and ligated into pSW177 [42]. pGT0285 was generated by amplifying the luminal domain with primers B29 containing a BamHI cut site and B30 that contain HA-HDEL-NcoI overhang sequences followed by ligation into BamHI/NcoI linearized pGT0223. The PGK promoter was amplified from pGT0121 with primer pair B36-37 and subsequently digested with NotI and BamHI and ligated with BamHI/Not1 linearized pGT0285 to make pGT0289. The plasmids BGT0261 and BGT0262 were generated by digesting pGT0223 and pGT0225 with NcoI/NotI and ligation into NcoI/Not1 digested pGT101. The plasmid pGT0334 was generated by Phusion site-directed mutagenesis from pGT0289 with GTO311-312 and GTO313-314 as previously described [92]. pGT0442 and pGT0443 was generated from pGT0261 and pGT0262 respectively by Phusion site-directed mutagenesis using the primer pair HN107-108 as previously described [92]. pGT0448 was a gift from Madhusudan Dey and constructed as previously described [93]. pGT0557 and pGT0558 were generated from pGT0330 and pGT0201 respectively by Phusion site-directed mutagenesis using the primer pair HN107-108. The split Venus constructs pGT0544 and pGT0546 were generated by Gibson assembly to join HA-VN173 (synthesized by Gblock) with HN177-178 linearized pGT0330 and pGT0201. The split Venus constructs pGT0545 and pGT0547 were generated by Gibson assembly to join FLAG-VC155 (synthesized by Gblock) with HN177-178 linearized pGT0059 and pGT0435. Yeast knockout strains in a Δ*ire1* BY4741 background were constructed by homologous recombination with the following primers: Δhrd1 with HN187 and HNHN188, Δlcb4 with HN189 and HN190, Δsec22 with HN191 and HN192, Δscj1 with HN199 and HN200, Δtor1 with HN203 and HN204, Δopi3 with HN205 and HN206, Δste24 with HN207 and HN208, Δget1 withHN209 and HN210, Δpah1 with HN211 and HN212.

### Yeast genetic screen

The library used was the yeast deletion library [34]. Using the Synthetic Genetic Array (SGA) methodology [94], the reporter strains YGT1228 and YGT1202 were mated to the MAT*a* yeast deletion library containing a single gene deleted with Kan^R^. In short, following mating and sporulation on nitrogen starved medium plates for 7 days, the MAT*α* cells were ultimately passaged onto SD plates containing geneticin sulphate (200 µg/ml), hygromycin B (200 µg/ml) and the toxic amino acid derivatives canavanine (100 µg/ml) and thialysine (100 µg/ml) to select for strains carrying either Ire1/iΔLD and Kan-marked gene deletions. The genetic screen was condensed with the 384 Solid Pin Multi-Blot Replicator (V&P Scientific, San Diego, CA) and performed in 384 format until analysis. Cells were subsequently pinned from 384 spots on agar to four 96-well plates using the 96 Solid Pin Multi-Blot Replicator (V&P Scientific, San Diego, CA) and inoculated in 200 µl YPD medium per well and grown overnight at 30°C. An automated high-throughput sampler (HTS) connected to the LSRFortessa X-20 (BD, Franklin Lakes, NJ USA) was used to measure the relative levels of GFP and mCherry. The program FACSDiVA v 8.0 (BD, Franklin Lakes, NJ) was used to acquire data in .fcs file format. Files were read with the program FlowJo X 10.0.7r2 (FlowJo, LLC). GFP and mCherry were excited at 488 and 561 nm, collected through a 505 and 595 nm long-pass filters and a 530/30 and 610/20 band pass filter, respectively. Reporter GFP fluorescence levels were normalized to the constitutive TEF promoter driven mCherry expression to correct for non-specific GFP expression. The median readout from 10,000 cells was obtained. The log_2_ GFP/mCherry ratio of each mutant (m) was normalized to WT levels from each plate and used as final sample’s reporter level using Eq. 1.

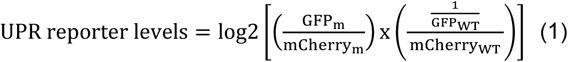

### *C. elegans* RNAi screen

To identify genes whose inactivation induced the UPR in *C. elegans*, we used a strain with a stably integrated *hsp-4p::gfp* reporter (strain SJ4005 *zcIs4 [hsp-4::GFP] V*), which is widely used as a reporter for UPR induction [10,12,95]. We compiled a list of 1695 metabolic genes from two different published datasets [39,40], and obtained RNAi clones for 1247 of these from the Ahringer RNAi library (Source BioScience). We performed the RNAi screen in duplicate in 12-well format in NGM-RNAi media seeded with appropriate HT115 RNAi bacteria. RNAi clones were tested in batches of 30-40 clones and each batch included negative (empty vector) and positive (*mdt-15* and/or *fat-6*) RNAi clones. Synchronized (by standard bleaching) L1 stage larvae were placed on RNAi bacteria lawns and allowed to develop into L4 stage larvae (~48 hours) and subsequently young adults (~72 hours); at both stages, *hsp-4p::gfp* levels were scored visually in an Leica M205FA upright fluorescent microscope. Clones causing visual developmental or growth delay were noted. Fluorescence was visually classified into three categories – low, medium, or high. We identified 118 RNAi clones that were scored as “medium” or “high” at either time point in both screens; these were subsequently validated in three small-scale repeat experiments, which yielded 40 *bona fide* hits. Hits were classified into strong, medium, or weak depending on their overall levels of induction across the population of worms examined. Sanger sequencing of the contained RNAi vectors revealed three clones with an insert other than the one identified in the library, and these were removed. Subsequently, the remaining 37 clones were tested for their ability to induce the *hsp-4p::gfp* reporter in strain SJ17 (*xbp-1(zc12) III; zcIs4 [hsp-4::GFP] V*). Choline supplementation was performed identically except that plates were additionally supplemented with 30 mM choline chloride. During choline supplementation and *xbp-1* dependency testing we observed that one clone, *ahcy-1*, only evoked very weak fluorescence and it was therefore excluded, leading to the final list of 36 genes (Table S2). *C. elegans* homologs of yeast hits were tested as above; of the 181 hits, 54 had *C. elegans* homologs, of which 38 had corresponding RNAi clones in the Ahringer RNAi library (Table S3).

### Indirect immunofluorescence

Indirect immunofluorescence was performed as previously described [96]. In brief, cells were grown to early log phase in selective media, fixed by 3.7% formaldehyde treatment and permeabilized. Monoclonal mouse anti-HA (1:500), anti-FLAG mouse M2 monoclonal antibody (1:500) and rabbit anti-Kar2 (1:1000) were used as primary antibodies. Mouse anti-Dylight 488 (1:500) and rabbit anti-Dylight 550 (1:500) were used as secondary antibodies. Samples were imaged with a Zeiss LSM 710 confocal microscope with a 100x 1.4 NA oil plan-Apochromat objective (Carl Zeiss MicroImaging). Images were analyzed using ImageJ 1.48v.

### Microscopy

For live-cell imaging, yeast cells were grown to an exponential phase at 30°C in 3ml of selective media. Samples were treated with 10 mM DTT for one hour or inositol depleted for 4 hours. Cells undergoing inositol depletion were washed 6 times before transferring to inositol free media. 500 µl of cells in selective media were placed on slides coated with 10 mg/ml Concanavalin A (Sigma-Aldrich, St. Louis, MO) mounted onto AttofluorTM cell chambers (Thermo Fisher, Waltham, MA) and imaged at room temperature. Samples were imaged with a Zeiss LSM 710 confocal microscope with a 100x 1.4 NA oil plan-Apochromat objective (Carl Zeiss MicroImaging). Images were analyzed using ImageJ 1.48v.

### Spotting growth assay

Cells were grown overnight in 3 ml of selective media at 30°C and diluted to 0.2 OD_600_/ml from which three 10-fold serial dilutions were prepared and spotted on selective plates (0.25 µg/ml tunicamycin or 1 mM choline were added to the plates when indicated). Plates were incubated at 30°C until the appearance of colonies.

### Alkaline carbonate extraction

Alkaline carbonate extraction was performed as previously described [97]. In brief, cells were grown to early log phase and the equivalent of 50 OD_600_ of cells were harvested. Cells were resuspended in 10 mM sodium phosphate buffer pH 7.0, 1mM PMSF and protease inhibitor cocktail (PIC) (Roche). An equal volume of 0.2 M sodium carbonate (pH 11.5) was added to cell lysates and incubated 30 minutes at 4°C and spun down at 100,000 × *g* for 30 min, 4°C. The pellet (membrane fraction) was solubilized in 3% SDS, 100 mM TrisCl, pH 7,4, 3 mM DTT and incubated at 95°C for 10 mins. Proteins from total cell lysate and supernatant fractions (collected from centrifuged lysate) were precipitated with 10% trichloroacetic acid (TCA) and spun down 30 min at 18,400 × *g*, 4°C. Proteins were resuspended in TCA resuspension buffer (1 mM Tris, pH 11, 3% SDS) and incubated 10 min at 95°C. Solubilized proteins were separated by SGS-PAGE and transferred to nitrocellulose for immunoblot analysis. Protein loading buffer was added to each fraction and separated by SDS-PAGE followed by immunoblot analysis.

### Immunoblot

Cells were grown to an early log phase overnight at 30°C. Tunicamycin was added to a final concentration of 2.5 µg/ml and incubated at 30°C for 1h, when indicated. Harvested cells were resuspended in 10% TCA was added to resuspended cells followed by the addition of 0.5 mm zirconium beads. Cells were disrupted by two 30 s cycles. The lysate was transferred to a new tube and combined with a 10% TAC bead wash. The precipitate was pelleted by centrifugation and vortexed in TCA resuspension buffer (100 mM Tris pH 11, 3% SDS, 1 mM PMSF). The samples were incubated 10 min at 95°C and spun down 15 min at 18,400 × *g*, 4°C. A portion of the extract was separated by SDS-PAGE using a 15% gel and transferred to nitrocellulose. The blots were probed with primary antibodies followed by secondary goat anti-mouse IgG-IRDye 800 (LI-COR Biosciences) and goat anti-rabbit IgG-IRDye 680 (LI-COR Biosciences, Lincoln, NE) antibodies. Membranes were washed in TBS and visualized with the Odyssey CLx imaging system (Li-COR).

### qPCR

Cells were grown to an early log phase overnight at 30°C. Tunicamycin was added to a final concentration of 2.5 µg/ml and incubated at 30°C for 1h, when indicated. Total RNA was extracted using RNeasy Mini Kit (Qiagen) following manufacture’s protocol. DNase treatment in columns was carried out with RNase-free DNase (Qiagen, Venlo, Netherlands) following the manufacturer’s protocol. cDNA was synthesized from 2 μg of total RNA using RevertAid reverse transcriptase (Thermo Fisher, Waltham, MA) following manufacturer’s protocol. SYBR Green qPCR experiments were performed following the manufacturer’s protocol using a QuantStudio 6 Flex Real-time PCR system (Applied Biosystems, Waltham, MA). cDNA (30 ng) and 50 nM of paired-primer mix were used for each reaction. Relative mRNA was determined with the comparative Ct method (ΔΔCt) normalized to housekeeping gene *ACT1*. Oligonucleotide primers used are listed in Table S8.

### β-galactosidase assay

The β-galactosidase assay was performed as previously described [83]. Typically, cells were grown to an early log phase overnight at 30°C. Tunicamycin was added to a final concentration of 2.5 µg/ml and incubated at 30°C for 1h, when indicated. Four OD_600_ of cells were collected and resuspended in 75 µl of Z buffer (125 mM sodium phosphate pH 7.0, 10 mM KCl, 1 mM MgSO_4_, 50 mM β-mercaptoethanol). An aliquot of 25 µl was transferred into 975 µl of ddH_2_O and the absorbance was measured at 600 nm. To the remaining resuspension, 50 µl chloroform and 20 µl 0.1% SDS was added and the resulting mixture was vortexed vigorously for 20 s. The reaction started with 700 µl of 2 mg/ml of o-nitrophenyl-β-D-galactoside (ONPG, Sigma) in Z buffer. The reaction was quenched with 500 µl of 1 M Na_2_CO_3_, and the total reaction time was recorded. Samples were spun for 1 min at maximum speed. Absorbance of the resulting supernatant was measured at 420 nm and 550 nm. The β-galactosidase activity was calculated using Eq. (2).

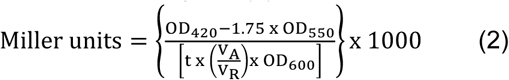

The values were then normalized to the activity of WT.

### Chromatin immunoprecipitation

Chromatin Immunoprecipitation (ChIP) was performed as previously described [98–100]. Typically, cells were grown to an early log phase. Tunicamycin was added to a final concentration of 2.5 µg/ml and incubated at 30°C for 1 h, when indicated. Forty OD_600_ of cells were collected, resuspended in 35 ml of selective, and fixed 20 min with 3.7% formaldehyde at 25°C. The reaction was quenched by adding 400 mM glycine. After an incubation of 5 min, cells were washed once with ice-cold TBS and resuspended in SDS lysis buffer (50 mM TrisCl pH 8.0, 10 mM EDTA, 1 % SDS). Cell lysates from these samples were sonicated for 8 cycles 10 s of 30% amplitude (Precellys 24, Bertin Instruments), with 50 s incubation on ice between intervals. Samples were diluted with the ChIP buffer (16.7 mM TrisCl pH 8.0, 167 mM NaCl, 1.1% Triton X-100, 0.01% SDS) to obtain a final concentration of 0.1% SDS. Forty microliters of protein G/salmon sperm DNA agarose beads and anti-HA in 1:500 dilution (Covance) were added followed by overnight incubation at 4°C. Beads were washed twice with low salt wash buffer (20 mM TrisCl, pH 8.0, 150 mM NaCl, 2 mM EDTA, 1% Triton X-100, 0.1% SDS) and high salt wash buffer (20mM TrisCl pH 8.0, 500 mM NaCl, 2 mM EDTA, 1% Triton X-100, 0.1% SDS), once with LiCl wash buffer [10 mM TrisCl, pH 8.0, 1 mM EDTA, 0.25M LiCl, 1% IGEPAL-CA630 (Sigma-Aldrich), 1% deoxycholic acid], twice with TE buffer (10 mM TrisCl, pH 8.0, 1 mM EDTA). Bound Hac1-HA was eluted by incubating the beads 20 min with 250 µl elution buffer (1% SDS, 0.1 M NaHCO_3_) at 30°C and repeated once. NaCl was added to the combined elution to a final concentration of 0.3M and incubated overnight at 65°C. Released DNA fragments were purified using the QiAprep Spin Miniprep Kit according to manufacturer’s protocol. Specific primers were designed approximately 75 bases up and downstream of the predicted UPRE motif (Table S8). The input DNA was diluted 100 times and ChIP DNA was pre-amplified 8 times with a primer mix prior to quantitative real-time PCR (qPCR). Quantitative PCR was performed following the manufacturer’s protocol using a CFX Connect Real-Time PCR Detection System (Bio-Rad). The Ct value obtained was used to calculate fold enrichment between experimental sample and normalized input using Eq. 3 and 4.

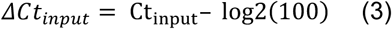

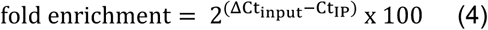

Results are representative of three biological replicates. *P* values were calculated using one-tailed Student’s t test.

### DNA microarray

Cultures were grown to an OD_600_/ml of 0.25 at 30°C in selective synthetic complete media. The UPR was induced in WT cells by 1 h incubation of 2 mM DTT when indicated. Cells were harvested from cultures at cell density of less than 0.5 OD_600_/ml. Total RNA was extracted by the hot acid phenol method as previously described [101]. Total RNA was subsequently cleaned up using the RNeasy Mini Kit (Qiagen). RNA quality control was carried out using the Agilent RNA Nano 6000 Chip (Agilent Technologies). RNA was prepared from independent triplicate samples. Probe preparation and microarray construction and analysis were performed as previously described [101–103]. Probes were prepared using the Low Input Quick Amp Labeling System with 100 ng of Total RNA as starting material following manufacturer’s instructions, which included One-Color Microarray-Based Gene Expression Analysis Protocol Version 6.5 (Agilent Technologies), and were hybridized on a Custom Microarray Agilent GE 8×60K array. Arrays were scanned using a high-resolution DNA Microarray Scanner, model G2505C (Agilent Technologies). Data were analyzed using GeneSpring GX software (Agilent Technologies). Differentially expressed genes were deemed significant with fold-change > 1.5 and ANOVA *P* values < 0.05. UpSetR was used to compare and visualize set intersections of significantly upregulated genes in a matrix-style layout [104]. GO terms analysis from gene lists acquired from the intersections was performed with DAVID [105]. Heat map in the figure was generated using R Studio. The DNA microarray data discussed in this publication has been deposited in NCBI’s Gene Expression Onmibus (GEO) under series number GSE131146.

### Cell labeling and immunoprecipitation

Cell labeling and immunoprecipitation was carried out as previously described [48]. Typically, 3 OD_600_ units of early log phase cells were labeled with 80 mCi of L-[^35^S]-methionine/cysteine mix (Perkin Elmer). Immunoprecipitated proteins were separated on SDS-PAGE and exposed to phosphor screens. Exposed screens were visualized using a Typhoon 8600 scanner (GE Healthcare).

### Statistics

The error bars denote standard error of the mean (SEM), derived from at least three biological replicates, unless otherwise indicated. *P* values were calculated using two-tailed Student’s t test unless otherwise indicated in the figure legends and reported as values in figures. Scatter plots were plotted using Graphpad Prism 8.

## AUTHOR CONTRIBUTIONS

Conceptualization: G.T.; Methodology: N.H., H.W., J.X., S.T., G.T.; Formal analysis: N.H, H.W., J.X., J.H.K.; Investigation: N.H., H.W., J.X., B.G., J.H.K.; Resources: N.H., H.W., J.X. ; Writing - original draft: N.H., H.W., J.X., S.T., G.T.; Writing - review & editing: N.H., H.W., J.X., S.T., G.T.; Supervision: S.T., G.T.; Project administration: G.T.; Funding acquisition: N.H., J.H.K., J.X., S.T., G.T.

## ACKNOWLEDGEMENTS

We are grateful to Dr. Maya Schuldiner for providing reagents and technical support to carry out our genetic screen. We thank Dr. Bhawana George for optimizing pilot experiments and members of Thibault lab for critical reading of the manuscript. This work was supported by the Nanyang Assistant Professorship program from the Nanyang Technological University (G.T.), the National Research Foundation, Singapore, under its NRF-NSFC joint research grant call (NRF2018NRFNSFC003SB-006 to G.T.), the Nanyang Technological University Research Scholarship to N.H., J.H.K. (predoctoral fellowship, the Natural Sciences and Engineering Research Council of Canada Discovery grant (RGPIN-2018-05133 to S.T.), and BCCHR Canucks for Kids Graduate and UBC Affiliate Studentships to J.X. Some strains were provided by the CGC, which is funded by NIH Office of Research Infrastructure Programs (P40 OD010440).

