## Supplemental Information for "ER stress sensor Ire1 deploys a divergent transcriptional program in response to lipid bilayer stress"

### SUPPLEMENTAL FIGURES

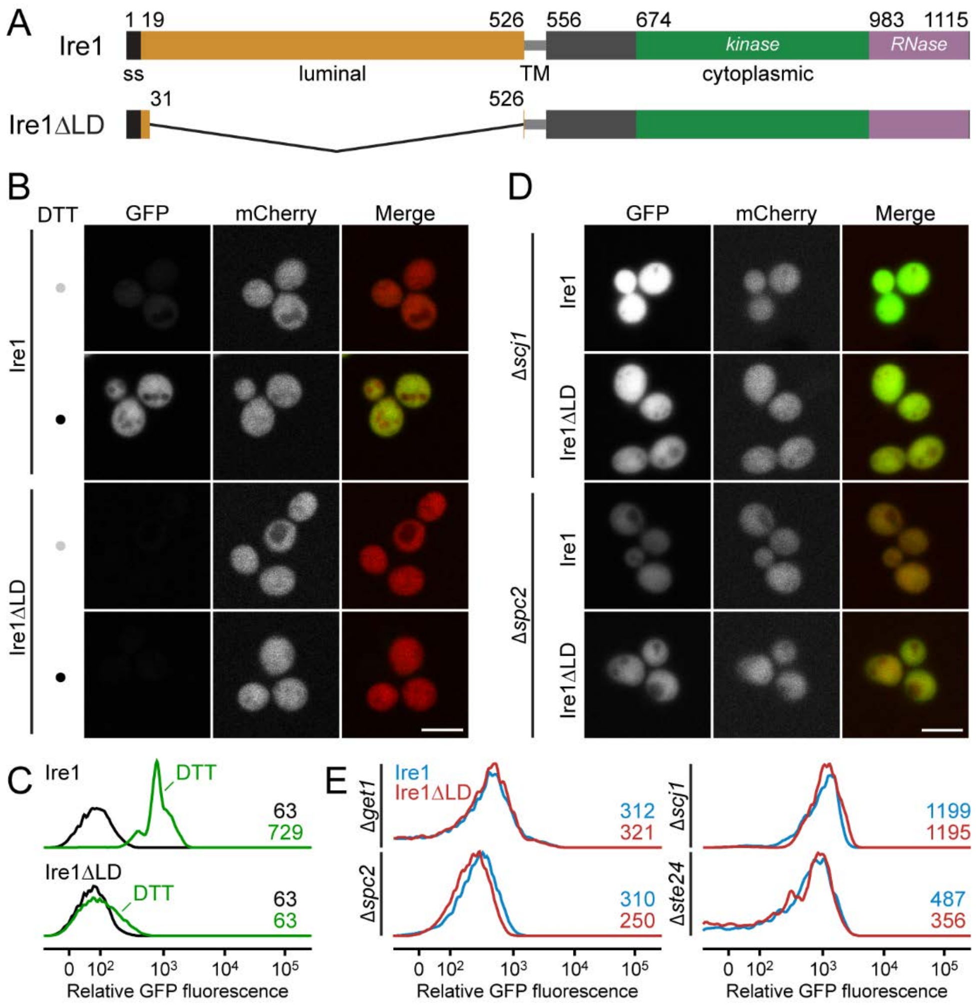


**Figure S1** (related to Figure 1)**. The UPR is activated in *IRE1* and *IRE1*Δ*LD* cells.**

(A) Schematic representation of Ire1 and Ire1ΔLD domain boundaries.

(B) Query strains *IRE1* and *IRE1*Δ*LD* with genomically integrated *UPREpr-GFP* and *TEF2pr-mCherry* were incubated 1h with 10 mM DTT, when indicated, before visualization by confocal microscopy. Scale bar, 5 µm.

(C) Flow cytometric histograms of strains and conditions as in (A). Numbers represent the maximum relative GFP fluorescence.

(D) Query strains with mutation Δ*scj1* or Δ*spc2* were visualized as in (A).

(E) Flow cytometric histograms of *IRE1* or *IRE1*Δ*LD* query strains with mutation Δ*get1*, Δ*spc2,* Δ*scj1* and Δ*ste24*. Numbers represent the maximum relative GFP fluorescence.

Images shown are representatives of three independent experiments.

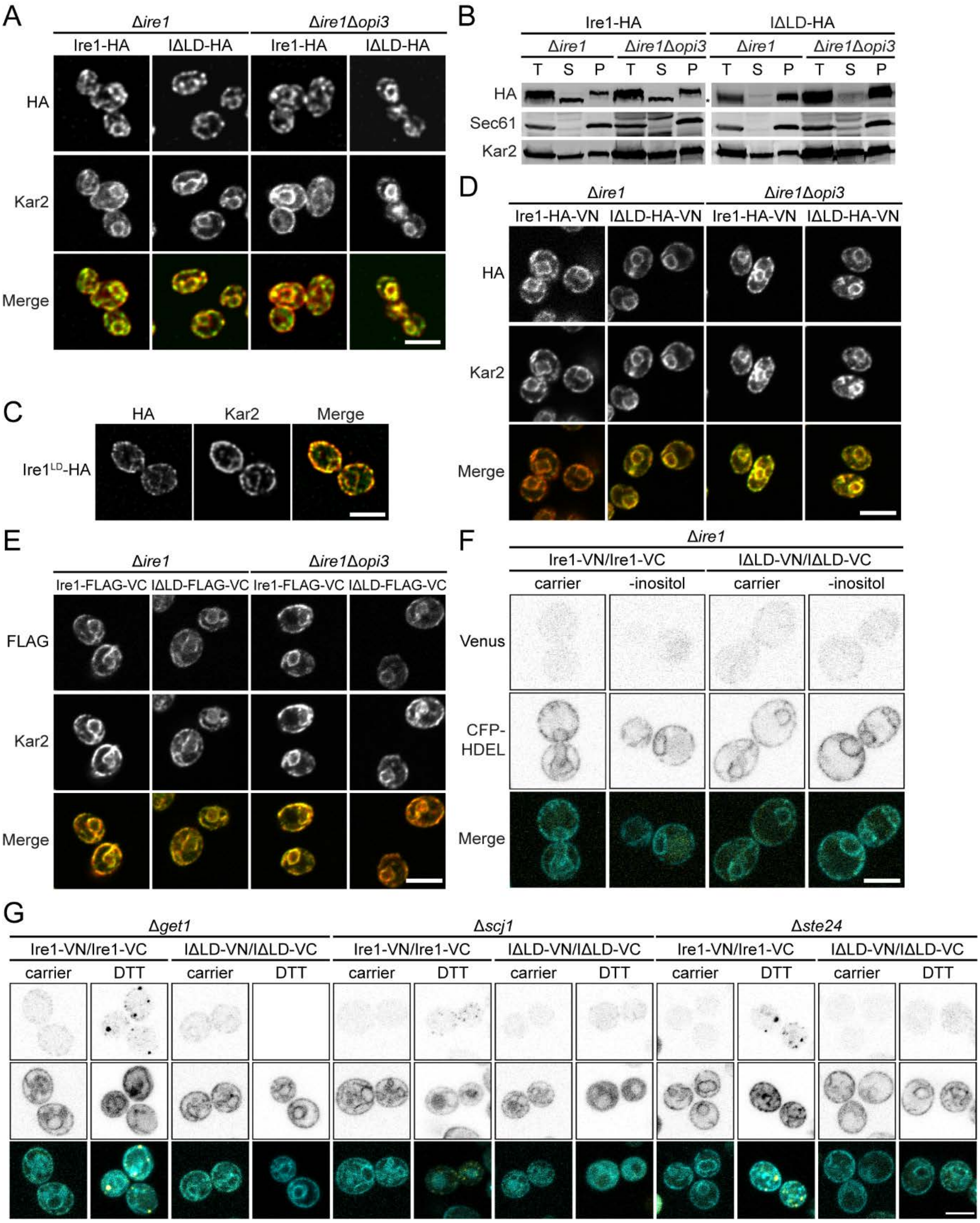


**Figure S2** (related to Figures 3 and 4)**. Ire1 and Ire1ΔLD are properly integrated into the ER membrane.**

(A) Cells were grown to early log phase in selective synthetic complete media before being fixed in formaldehyde and permeabilised. Staining was performed using anti-HA and anti-Kar2 primary antibodies. Scale bar, 5 µm.

(B) Membrane prepared from the indicated cells were treated with 0.1 M sodium carbonate, pH 11, for 30 min on ice. A portion was kept as the total fraction (T), and the remaining was subjected to centrifugation at 100,000 X *g*. Supernatant (S) and membrane pellet (P) fractions were collected and analyzed by immunoblotting. Proteins were detected using antibodies against HA. Kar2 and Sec61 serve as soluble and integral membrane protein controls, respectively.

(C-D) Cells were grown and treated as in (A). Scale bar, 5 µm.

(E) Cells were grown and treated as in (A). Staining was performed using anti-FLAG and anti-Kar2 primary antibodies. Scale bar, 5 µm.

(F) Cells co-expressing the pair of split Venus fragments to monitor *IRE1-HA-VN173* and *IRE1-FLAG-VC155* or *IRE1ΔLD-HA-VN173* and *IRE1ΔLD-FLAG-VC155* to monitor dimerization *in vivo* by bimolecular fluorescence complementation assay (BiFC) in Δ*ire1* and Δ*ire1*Δ*opi3*. Cells were depleted of inositol (-inositol). Cells also expressed the CFP-HDEL ER marker. Scale bar, 5 µm.

Images shown are representatives of three independent experiments.

(G) Mutant strains Δ*get1*, Δ*scj1*, and Δ*ste24* treated as in (F).

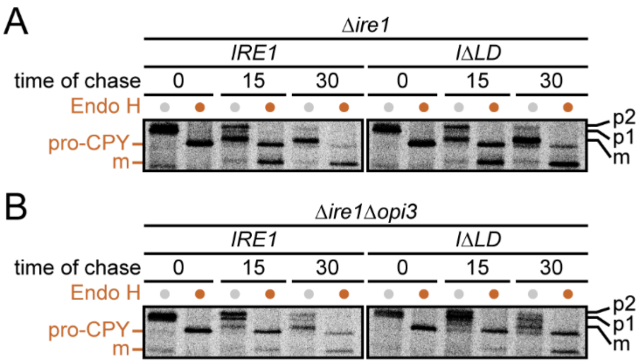


**Figure S3** (related to Figure 3)**. *IRE1$^{LD}$* is dispensable for CPY biogenesis in Δ*opi3* cells.**

Cells were grown to early log phase in selective synthetic complete media before being pulse-labeled with L-[$^{35}$S]-methionine/cysteine for 5 min followed by a chase at the indicated time. Immunoprecipitated protein using anti-CPY were treated with Endo H, when indicated, and resolved by SDS-PAGE and visualized by phosphoimager analysis. Images shown are representatives of three independent experiments.

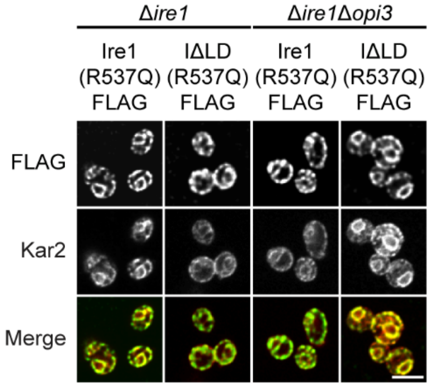


**Figure S4** (related to Figure 5)**. Ire1 and Ire1ΔLD containing the transmembrane mutation R537Q are properly localized into the ER membrane.**

Cells were grown to early log phase in selective synthetic complete media before being fixed in formaldehyde and permeabilised. Staining was performed using anti-FLAG and anti-Kar2 primary antibodies. Cells were incubated 1h with 10 mM DTT, when indicated. Scale bar, 5 µm. Images shown are representatives of three independent experiments.

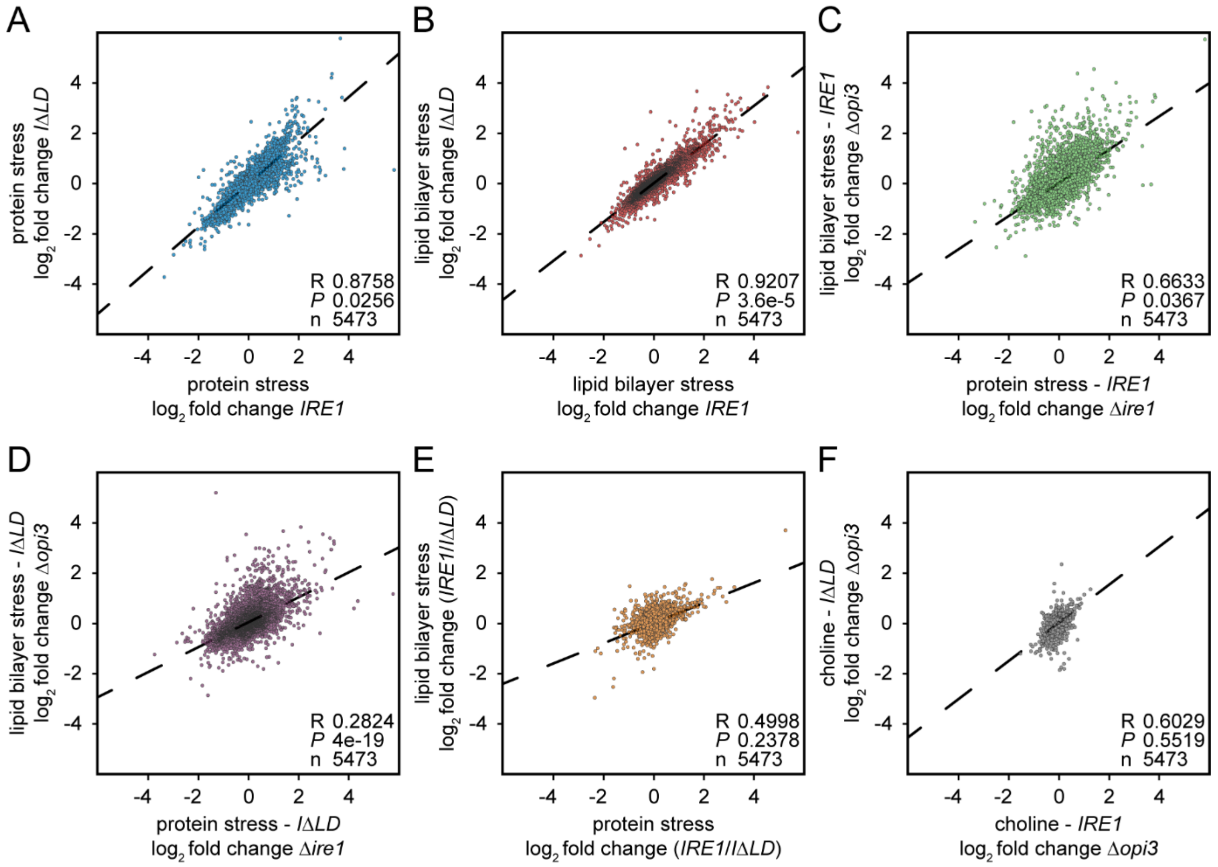


**Figure S5** (related to Figure 6)**. Transcriptional changes reveal LD-specific strategies in coping with different stresses.**

(A) Pearson correlation of relative RNA abundance in *IRE1* and *IRE1ΔLD* treated with 2 mM DTT.

(B) Pearson correlation of relative RNA abundance in *IRE1Δopi3* and *IRE1ΔLDΔopi3*.

(C) Pearson correlation of relative RNA abundance in *IRE1* treated with 2 mM DTT and *IRE1Δopi3*.

(D) Pearson correlation of relative RNA abundance in *IRE1ΔLD* treated with 2 mM DTT and *IRE1ΔLDΔopi3*.

(E) Pearson correlation of relative RNA abundance ratio of *IRE1*/*IRE1ΔLD* treated with 2 mM DTT and *IRE1Δopi3*/*IRE1ΔLDΔopi3*.

(F) Pearson correlation of relative RNA abundance in *IRE1Δopi3* and *IRE1ΔLDΔopi3* supplemented with 1 mM choline.

**Table S6** (related to Figure 1-6)**. Yeast strains used in this study**

| Strain | Genotype | Source |
|---|---|---|
| W303a | *leu2-3,112 trp1-1 can1-100 ura3-1 ade2-1 his3-11,15* | (Cox et al., 1993) |
| BY4741 | *his3Δ1 leu2Δ0 met15Δ0 ura3Δ0* | (Brachmann et al., 1998) |
| ESY248 | *MAT*a, *ire1::TRP1*, W303 background | (Spear and Ng, 2005) |
| YMS612 | *MAT*α, *UPRE-GFP::URA TEF2-cherry::Met his3Δ1 leu2Δ0 lys2+ met15Δ0 ura3Δ0 can1Δ::STE2pr-sp HIS5 lyp1Δ::STE3pr-LEU2 cyh2* BY4741 | (Cohen et al., 2017) |
| YGT0268 | *MAT*a, *ire1:: KANMX*, *opi3:: KANMX*, W303 background | This study |
| YGT0271 | *MAT*a, *ire1:: KANMX*, W303 background | This study |
| YGT0705 | *MAT*a, *ire1::TRP1*, *opi3:: KANMX*, W303 background | (Thibault et al., 2012) |
| YGT1202 | *MAT*α, *ire1::IRE1ΔLD::Hyg*$^{R}$ *UPRE-GFP::URA TEF2-cherry::Met* YMS612 background | This study |
| YGT1228 | *MAT*α, *ire1::IRE1::Hyg*$^{R}$ *UPRE-GFP::URA TEF2-cherry::Met* MS612 background | This study |

**Table S7** (related to Figure 3-6)**. Plasmids used in this study**

| Plasmid | Encoded protein | Promoter | Vector | Source |
|---|---|---|---|---|
| pGT0012 | β-galactosidase | *UPRE-CYC1* | pRS314 | (Cox and Walter, 1996) |
| pGT0059 | Ire1 | *IRE1* | pRS315 | This study |
| pGT0201 | Ire1ΔLD | *IRE1* | pRS313 | This study |
| pGT0223 | Ire1 | *PRC1* | pRS315 | This study |
| pGT0224 | Ire1ΔLD | *PRC1* | pRS315 | This study |
| pGT0262 | Ire1-FLAG | *PRC1* | pRS313 | This study |
| pGT0263 | Ire1ΔLD-FLAG | *PRC1* | pRS313 | This study |
| pGT0289 | Ire1 LD-HA | *PGK1* | pRS315 | This study |
| pGT0330 | Ire1 | *IRE1* | pRS313 | This study |
| pGT0421 | Kar2 | *TDH3* | pMR366 | (Thibault et al., 2012) |
| pGT0435 | Ire1ΔLD | *IRE1* | pRS315 | This study |
| pGT0442 | Ire1(R537Q)-FLAG | *PRC1* | pRS313 | This study |
| pGT0443 | Ire1ΔLD(R537Q)-FLAG | *PRC1* | pRS313 | This study |
| pGT0448 | 3X-HA-Hac1 | *HAC1* | pRS316 | (Sathe et al., 2015) |
| pGT0544 | Ire1-HA-VN | *IRE1* | pRS313 | This study |
| pGT0545 | Ire1-FLAG-VC | *IRE1* | pRS315 | This study |
| pGT0546 | Ire1ΔLD-HA-VN | *IRE1* | pRS313 | This study |
| pGT0547 | Ire1ΔLD-FLAG-VC | *IRE1* | pRS315 | This study |
| pGT0557 | Ire1(R537Q) | *IRE1* | pRS313 | This study |
| pGT0558 | Ire1ΔLD(R537Q) | *IRE1* | pRS313 | This study |
| pMJ0012 | mCherry | *TEF2* | pFA6-natMX6 | (Jonikas et al., 2009) |
| pMS0001 | GFP | *UPRE-CYC1* | pRS306 | (Jonikas et al., 2009) |

**Table S8** (related to Figure 1-6)**. Oligonucleotide primers used in this study**

| Primer | Sequence (5' to 3') |
|---|---|
| HN46 | CTGGAAATTTCACCCGGC |
| HN47 | CAGTAACACGTTAATCTTTAGAATTAC |
| HN56 | CAGCCCTTATTTTTTTTCCAAG |
| HN57 | GGGATGCGTGGTATTG |
| HN58 | CACGTACATTGCTGTGG |
| HN59 | AGTGCACGCTCAACT |
| HN60 | CTCCTGATAGACAATGAAATATACC |
| HN61 | AGTGCTACTAGGCGATG |
| HN62 | CCCTATCTTTTTTTTTTTCTCGC |
| HN63 | ATGCAGTGAAACTAAGGATAC |
| HN64 | GCAAGATGATGTTCAACCT |
| HN65 | TACTTGGCATATTGCTTAACTC |
| HN66 | GTGAAAAATGCATCCAAAAGA |
| HN67 | TACTACATTCGGAAATGGTTG |
| HN107 | CCAGTCTCTATAATTTGATATACTAGACTTCC |
| HN108 | AGTATTTCTGTTGTTATTTCTCATTTTTTGTG |
| HN117 | TTAGGCGGCCGGAGCGTAA |
| HN118 | GGGCTGCAGGAATTCGATATCAAGCT |
| HN119 | GACATTTAATTTTATAAATACATATCTCATAAACAGCGACATGGAGGCCCAG |
| HN120 | AACTGGAGTAGTATGTCGATGTTCGATTATAAATAACGTTCTTAATACTAAC |
| HN121 | GTTTATGAGATATGTATTTATAAAATTAAATGTCTGTC |
| HN122 | ATCGAACATCGACATACTACTCC |
| HN148 | ACAAAGAAGTAATGAACTTAAATGCTATTATACAG |
| HN149 | CCGTCCCAAACATTGTCATAGATTC |
| HN153 | CATGTTCATGCCCCTCTGC |
| HN177 | TGAATACAAAAATTCACGTAAAATTTGATCGTCAC |
| HN178 | TAACATGTTCATGCCCCTCTGC |
| HN187 | TCAATTGCAATTTGTAAGAGAAGGGGAGAAAGACAAAATAATAATATGGGTAAAAAGCCT |
| HN188 | GTTTTTTTCTTTAAAAAAAACTATGTATAATATAAAACATGCAATTTATTCCTTTGCCCT |
| HN189 | CAAAAGTCTAGCAGCGAAAAGTACGCGAAGAATCTACTATAGATAATGGGTAAAAAGCC |
| HN190 | TCTTTTTACAAAAAAATCATTTTTGAAGGAAAATATAACGTTAATTTATTCCTTTGCCCT |
| HN191 | AACCCTGACAGTGACACCCCGTTACACACTCACAATTAAGTAGGGATGGGTAAAAAGCCT |
| HN192 | GACCAAATTGATCGGGATTTGTGATGTGGGATGATGGGGTGACGTTTATTCCTTTGCCCT |
| HN199 | ATAGGCCAGAAGAGCGTGCATTGGCTGGCGAAAAGATCGAGGACAATGGGTAAAAAGCCT |
| HN200 | GTTGTCTATCTATATATGCATGTGTGCGTACGTAGGATTATCTGTTTATTCCTTTGCCCT |
| HN203 | GGTAAAGTGAAACATACATCAACCGGCTAGCAGGTTTGCATTGATATGGGTAAAAAGCCT |
| HN204 | AAAATAAATAGTAAACAAAGCACGAAATGAAAAATGACACCGCAGTTATTCCTTTGCCCT |
| HN205 | CAATTGAAGACAACAAGAATAGCGCAAGTCAAGCGATGAAGGAGTATGGGTAAAAAGCCT |
| HN206 | ATAGCATAGGCTTCTAACATTATAGAATATATAGAAATAGAGCACTTATTCCTTTGCCCT |
| HN207 | GGAAGGAAAAAAAAGGAGGAAATAGAAAACTGCAGGCCTTTATTCATGGGTAAAAAGCCT |
| HN208 | CGAAGCTGAATTTAACGGTACATGCTAATATGTGTACTCTATAGATTATTCCTTTGCCCT |

| Primer | Sequence (5' to 3') |
|---|---|
| HN209 | AGTTTGCAATCCTTGAACTACGTCTAGTTGATTGAAATAGGAGAAATGGGTAAAAAGCCT |
| HN210 | TCCAATACATAAACATATTATATATACGTACATAATGTAATAACATTATTCCTTTGCCCT |
| HN211 | AACATACAGGGAAGAAATTACTGAAGATAGACACATCGGTCGATTATGGGTAAAAAGCCT |
| HN212 | TATGCAGTATGGATCGTTATAAATAATATTCGGCTACAAGAATCTTTATTCCTTTGCCCT |
| GTO95 | GACCAAGAGACTTCATGGGAC |
| GTO96 | GAATTCAAACCTGACTGCGC |
| GTO103 | GGTTGCTGCTTTGGTTATTGA |
| GTO104 | TTTTGACCCATACCGACCAT |
| GTO311 | 5'P ATGAAAAGGTCTACTGTGGATCGATGAGAA |
| GTO312 | 5'P CTTCAATTTTTCCAGAGTCATTAACC |
| GTO313 | ATATCAGCGTGCGGACCTGGT |
| GTO314 | AATTTCACCATTCAACATGTTTATAGTATACATTATAGTTCTC |
| GTO275 | ATTCCTCGAGAAACTCTGCTGCGCGCTGAA |
| GTO276 | GAATCTGCAGGCGAGAACGACAATGGGATTGAG |
| GTO277 | GAATCTGCAGAAACACACAAACGAGCAGTGTC |
| GTO278 | GAATGCGGCCGCAAGTCAGTGTTGAATAACTGGAGTAG |
| HWO15 | ACTCGGATCCATGCGTCTACTTCGAAGAAACAT |
| HWO16 | ACTGCCATGGCTGAATACAAAAATTCACGTAAAATTTGATCGT |
| B29 | CGCGGATCCATGCGTCTACTTCGAAGA |
| B30 | CATGGCGGCCGCTCACAATTCGTCGTGAGCGTAATCTGGAACATCATATGGGTAATTTTGGTTCTTTTCATCTAATTC |
| B36 | ATGCGCGGCCGCAGACGCGAATTTTTCGAAG |
| B37 | GCATGGATCCTGTTTTATATTTGTTGTAAAAAGTAGA |
